# Transcranial Ultrasound Stimulation in Anterior Cingulate Cortex Impairs Information Sampling and Learning in Loss Contexts

**DOI:** 10.1101/2021.08.04.455080

**Authors:** Kianoush Banaie Boroujeni, Michelle K Sigona, Robert Louie Treuting, Thomas J. Manuel, Charles F. Caskey, Thilo Womelsdorf

## Abstract

Neuronal subgroups in anterior cingulate cortex (ACC) and the anterior striatum (STR) encode the reward structure of a given environment. But whether or how this reward information is used to guide information sampling, optimize decision making, or motivate behavior in cognitively challenging situations has remained elusive. Here, we causally tested these scenarios by transiently disrupting ACC and STR of rhesus monkeys with transcranial ultrasound with a learning task that independently varied cognitive and motivational demands. We found that disrupting the ACC, but not the STR, prolonged information sampling and reduced learning efficiency whenever the motivational payoff was low. These impairments were most pronounced at high cognitive demands and based on an inability to use loss experiences to improve performance. These results provide causal evidence that the ACC is necessary for motivation, to overcome anticipated costs from negative (loss) outcomes, and for cognition, to enhance visual information sampling during adaptive behavior.

**HIGHLIGHTS:**

- Transcranial ultrasound stimulation of the anterior cingulate cortex disrupts learning after loss experience.
- The ultrasound-induced learning deficit is exacerbated at high cognitive load.
- The ultrasound-induced learning deficit is accompanied by inefficient fixational information sampling.
- Anterior cingulate cortex causally supports credit assignment of aversive outcomes to visual features.

## INTRODUCTION

It is well established that the ACC and the anterior striatum contribute to flexible learning (*1*–*4*). Widespread lesions of either structure can lead to non-adaptive behavior. When tasks require subjects to adjust choice strategies, lesions in the ACC cause subjects to shift away from a rewarding strategy even after obtaining reward for a choice (*5*), and reduce the ability to use error feedback to improve behavior (*6*–*8*). Lesions in the striatum likewise reduce the ability to use negative error feedback to adjust choice strategies, which can take on the form of perseveration of non-rewarded choices (*9*), or inconsistent switching to alternative options after errors (*10*). A common interpretation of these lesion effects is that both structures are necessary for integrating the outcome of recent choices to update the expected values of possible choice options. According to this view, ACC and the striatum keep track of the reward structure of the different features and choice options in a given task environment.

However, it has remained contentious how such a neural representation of the current reward structure affects adaptive behavior. One perspective suggests the ACC uses reward information to compute the overall value for the subject to continue with the current choice strategy or switch strategies when the value to continue making similar choices is lower than the value of choosing alternative options (*11, 12*). An important extension of this viewpoint suggests that the representations of values in the ACC and the striatum do not only trigger behavioral changes, but also guide attention and information sampling to maximize rewards and minimize punishments (*13*). According to this model, lesioning the ACC or the striatum will alter information sampling behavior and reduce the ability of subjects to estimate the value of choice options properly. However, so far these predictions have not been tested directly with lesions to the ACC or the striatum.

An alternative viewpoint emphasizes that the ACC-striatum network uses reward information in conjunction with information about task demands to control how much effort and cognitive control will be exerted to succeed with a task (*4, 14*). Similar to the previous model, this theory suggests that neural circuits within the ACC uses information about recent reward- and punishment-outcomes to update the expected gains and losses of different choice options. In contrast to the previous model, however, the model suggests that the information about expected gains and losses are combined with an estimate of the cognitive demands or difficulty of a task in order to determine how valuable it is for the subject to put more effort into controlling cognitive performance (*15*). A key prediction of this model is that the ACC will be necessary to put effort into task performance when either the cognitive load increases, or when the expected payoff decreases and motivational challenges need to be overcome (*15*). These predictions still await to be causally tested with a task paradigm that disentangles the influence of cognitive demands and varying payoff.

Here, we set up an experiment that allowed us to test the predictions of both theoretical viewpoints using a transcranial ultrasound stimulation protocol that temporarily disrupted either the ACC or the striatum in rhesus monkeys performing a complex reward learning task. The task independently varied attentional load and the demands for information sampling from varying the payoff that subjects could receive for successful task performance (**Fig. 1A**,**B**). Subjects had to learn a feature-reward rule by choosing one of three objects that varied in features of only one dimension (low attention load, e.g., different shapes), or in features of two or three dimensions (high attention load, e.g., varying shapes, surface patterns, and arm types) (**Fig. 1C**). Independent of attentional load, we varied how motivationally challenging task performance was by altering whether the learning context was a pure *gain-only* context or a mixed *gain-loss* context. In the *gain-only contexts*, subjects received three tokens for correct choices, while in the *gain-loss contexts*, subjects received two tokens for correct choices and lost one already attained token when choosing objects with non-rewarded features (**Fig. 1A-C**). The loss of tokens triggers a vigilance response (*16*) and imposes a motivational conflict (Banaie Boroujeni et al., 2020). The task required monkeys to collect five visual tokens before they were cashed out for fluid rewards.

**Fig. 1.**
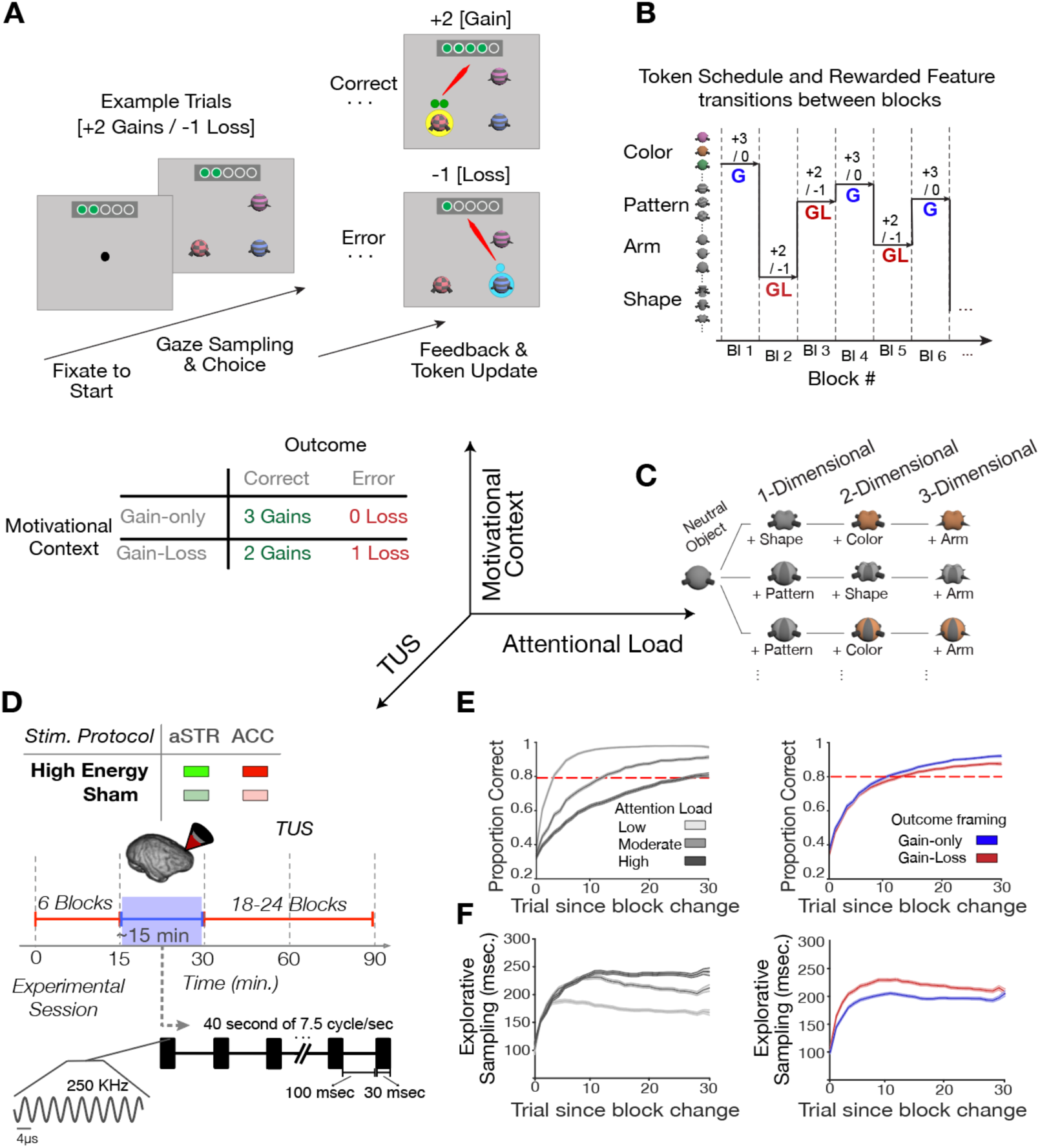
Task paradigm and TUS protocol. **A**, A trial started with central gaze fixation and appearance of 3 objects. Monkeys could then explore objects and choose one object by fixating it for 700 ms. A correct choice triggered visual feedback (a yellow halo of the chosen object) and the appearance of green circles (tokens for reward) above the chosen object. The tokens were then animated and traveled to a token bar on the top of the screen. Incorrect choices triggered a blue halo and in the gain-loss condition one blue token was shown that traveled to the token bar where one already attained token was removed. When ≥5 tokens were collected in the token bar, fluid reward was delivered, and the token bar reset to zero. **B**, In successive learning blocks, different visual features were associated with reward and blocks alternated randomly the gain-only (G) and gain-loss (GL) conditions. **C**, Attentional load varied by increasing the number of object features that varied from block to block from 1, 2 and 3. **D**, In each sonication or sham session the experiment was paused after 6 learning blocks. There are four experimental conditions; high energy TUS in ACC (H-ACC; red), or anterior striatum (H-aSTR; green), or sham ACC (S-ACC; dimmed red), or sham anterior striatum (S-aSTR; dimmed green). 30-ms bursts of transcranial ultrasound stimulation were delivered every 100-ms over a duration of 80-second (40-second each hemisphere). **E**, Proportion of correct choices over trials since block begin for different attentional loads (left panel; 1-3D, light to dark grey) and learning contexts (gain-only, blue; gain-loss, red). **F**, The average fixation duration of objects during explorative sampling prior to choosing an object in the same format as in (**E**). The lines show the mean and the shaded error bars are standard error of the mean.

With this design we found that disrupting the ACC, but not the anterior striatum with transcranial ultrasound stimulation, caused a learning deficit when subjects experienced low payoff (losses). This loss-triggered deficit was accompanied by inefficient information sampling and most pronounced at high attention load.

## RESULTS

We applied transcranial focused ultrasound (TUS) to disrupt the ACC (area 24) or anterior striatum (aSTR, head of the caudate nucleus) in two monkeys in separate learning sessions by adopting the same TUS protocol as in (*18, 19*). The sonication protocol imposed a ∼6mm wide / 40mm tall sonication region that has been shown previously to alter behavior in foraging tasks (*20*), to reduce functional connectivity of the sonicated area in macaques (*18*), and in in-vitro preparations to modulate neuronal excitability to external inputs (*21*). We provide detailed acoustic simulations of the ultrasound pressure dispersion around the target brain areas, the anatomical sub-millimeter targeting precision of TUS, and the validation of the applied ultrasound power through real-time power monitoring during the experiment in **Fig. S3**. We bilaterally sonicated or sham-sonicated the ACC or the aSTR in individual sessions immediately after monkeys had completed the first six learning blocks. Following the sonication procedure, monkeys resumed the task and proceeded on average for 23.6 (±4 SE) learning blocks (monkey W: 20.5±4; monkey I: 26.5±4) (**Fig. 1D**).

Across learning blocks, monkeys reached the learning criterion (≥80% correct choices over 12 trials) systematically later in blocks with high attentional load (LMEs, p<0.001; **Fig. 1E, Fig. S1A-C**). Both monkeys also showed longer foveation durations onto the objects prior to making a choice when the attentional load was high (LMEs, p<0.001; **Fig. 1F**; **Fig. S1D-E**). These longer foveation durations index more extensive explorative information sampling of object features at higher attentional load. Monkeys also learned slower and showed reduced plateau performance (**Fig. S2A-C**) as well as increased explorative sampling in blocks with gains and losses (gain-loss contexts) compared to blocks with only gains (gain-only contexts) (**Fig. 1F, Fig. S2D-E**; LMEs, p<0.001).

TUS in ACC but not sham-TUS in ACC or TUS or sham-TUS in the striatum (**Fig. 2A**) changed this behavioral pattern. ACC-TUS selectively slowed learning in the gain-loss contexts compared to the gain-only contexts (LME’s, t=2.67, p=0.007)) (**Fig. 2B**; **Fig. S4A**,**B**, **Supplementary Table 1**). ACC-TUS increased the number of trials needed to reach the learning criterion of 80% performance to 14.7±0.8 trials (monkey W/I: 14.3±1.2 / 15±1.2) relative to pre-TUS baseline (trials to criterion: 10.7±1.4; monkey W/I: 9.9±2.3 / 11.4±1.8) (Wilcoxon test, p=0.049; **Fig. 2B**; **Fig. S4C**,**D**). The learning speed with ACC-TUS in the gain-loss context was significantly slower to other TUS conditions (Kruskal-Wallis test, p=0.003), ACC-sham (pairwise Wilcoxon test, FDR multiple comparison corrected for dependent samples, p=0.019), and to the conditions aSTR-TUS and aSTR-sham (pairwise Wilcoxon test, FDR multiple comparison corrected for dependent samples, p=0.019, p=0.003) (**Fig. 2B**). This effect interacted with attentional load. The slower learning after ACC-TUS in the gain/loss condition was stronger when attentional load was intermediate (2 distracting feature dimensions) and high (3 distracting feature dimensions) (random permutation, p<0.05; LME’s, t=-2.8, p=0.004) (**Fig. 3C**,**D**; **Fig. S5A, Supplementary Table 2**). TUS did not affect learning in the gain-only contexts even at high attentional load (Kruskal-Wallis test, p=0.933) (**Fig. 3C**,**D**; **Fig. S5B**).

**Fig. 2.**
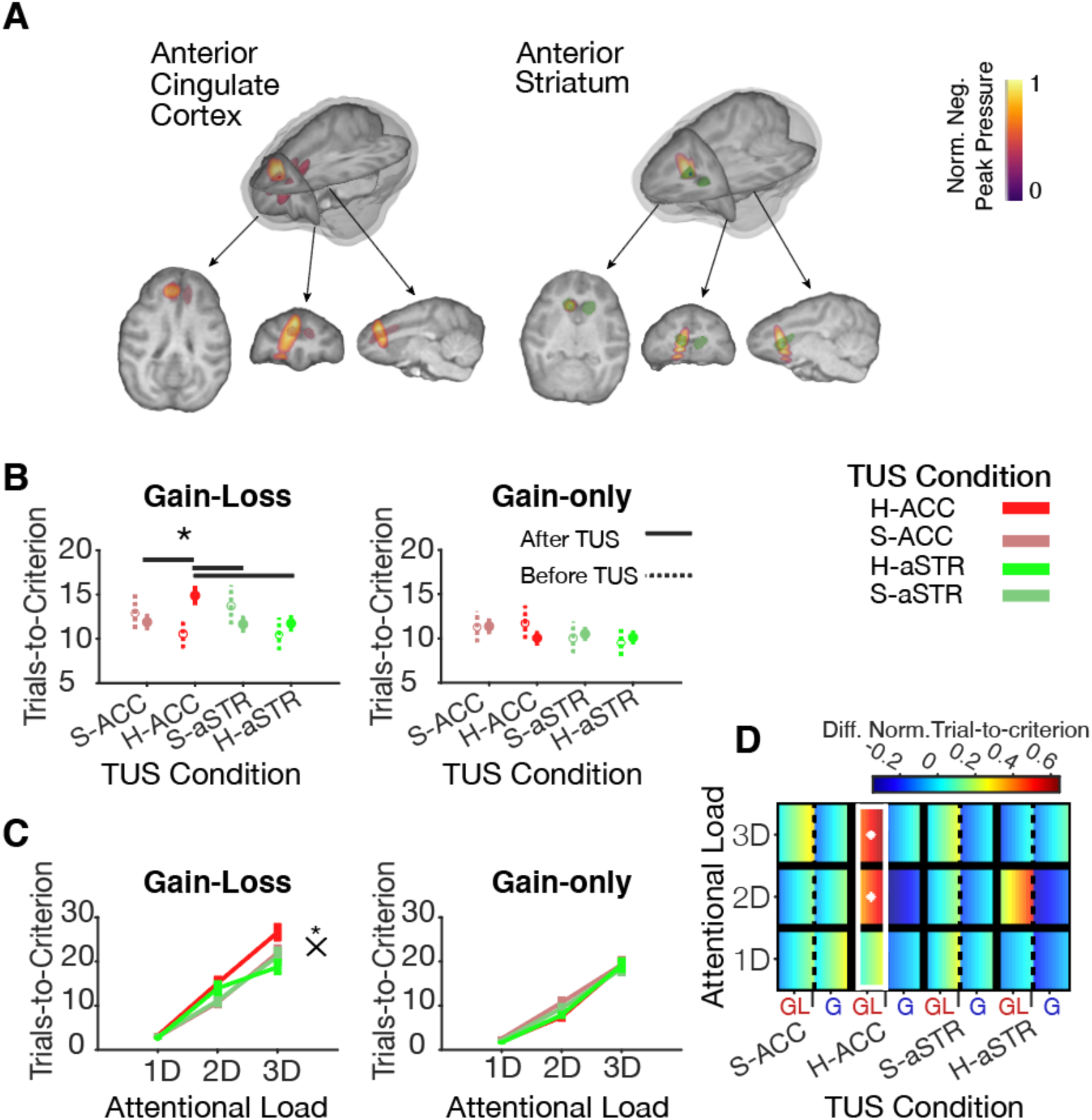
Transcranial ultrasound stimulation (TUS) of ACC and anterior striatum. **A**. The maximum negative peak pressure of TUS in an example session of one monkey shows the focus is within ACC (left) and anterior striatum (right). **B**. Learning is slowed down (trials-to-criterion increased) after TUS in ACC in the gain-loss context (left, LME’s, p=0.007) but not in the gain-only contexts (right, n.s.). Data represent means and the standard error of the mean. **C**. Learning is slowed with higher attentional load with a significant interaction in the gain-loss context (left, LME’s, p=0.007) but not in the gain-only context (right, n.s., for the full multiple comparison corrected statistical results see **Supplementary Table 1**,**2**). **D**. Marginally normalized trials-to-criterion is significantly higher with ACC-TUS in the gain-loss (GL) learning context at higher (2D and 3D) attentional load (random permutation p<0.05). Each cell is color coded with the mean value ± standard error of the mean with a low to high value gradient from left to right. The white rectangle shows that learning in that TUS condition (*x-axis*) is different from other TUS conditions. White asterisks indicate that an attentional load condition in a learning context in a TUS condition is significantly different from other TUS conditions.

**Fig. 3.**
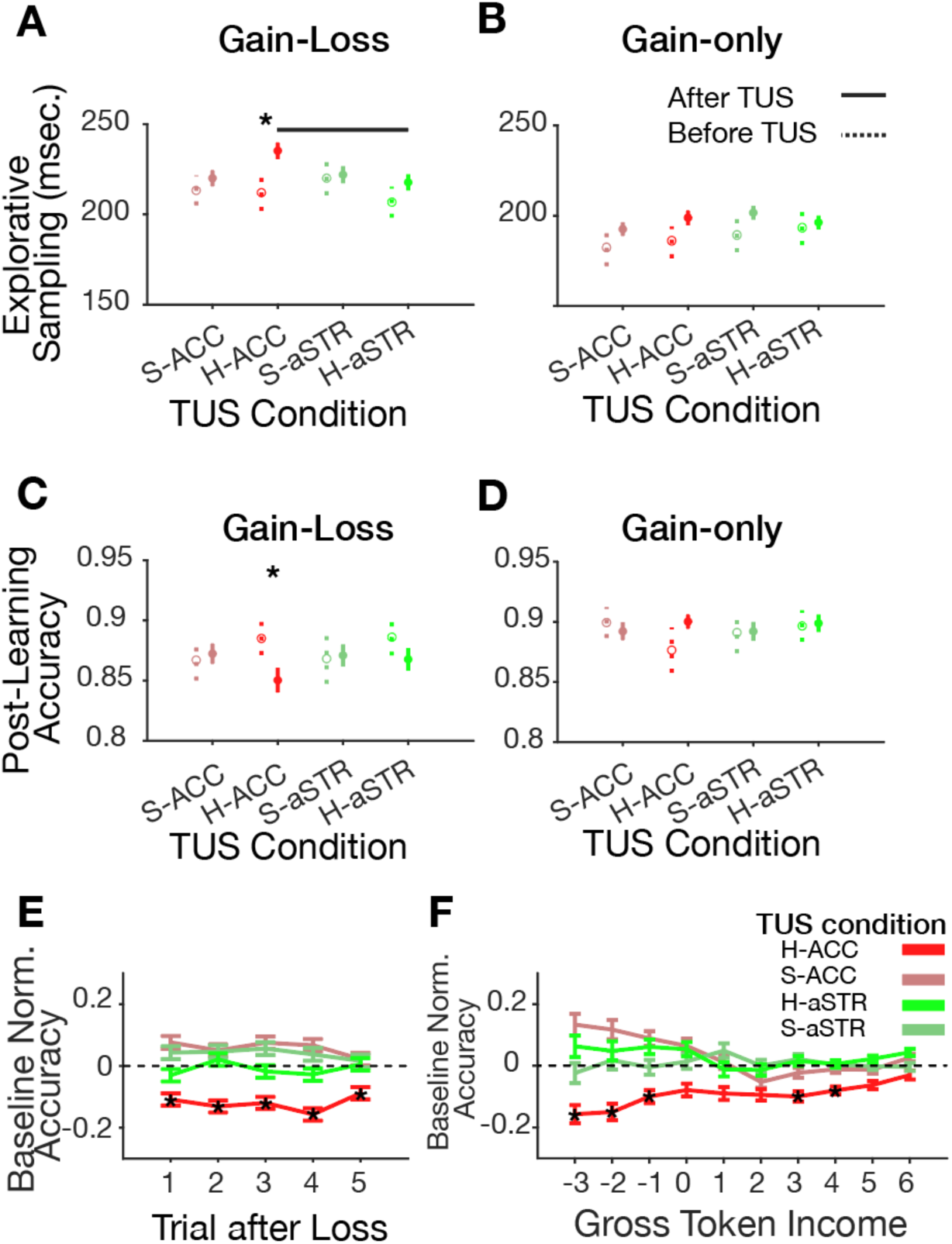
Transcranial ultrasound stimulation effect on explorative behavior and post-error adjustment. **A**,**B**. Explorative sampling (the duration of fixating objects prior to choosing an object) is increased with TUS in ACC in gain-loss (**A**) but not gain-only (**B**) learning context. **C**,**D**. TUS in ACC reduced post-learning plateau accuracy in gain-loss contexts (**C**) but not in gain-only contexts (**D**) (LMEs, p=0.04) (see **Supplementary Table 2**). With FDR correction, this accuracy effect was significant at the session level, but not at the block level, indicating a low effect size (**Fig. S8E** and **Supplementary Table 1**). **E**. The accuracy was overall reduced with TUS in ACC in the five trials after experiencing a token loss in the gain-loss context (random permutation, p<0.05). **F**. Gross token income (GTI) (*x-axis*) measures the signed average of tokens gained and lost in the preceding trials. TUS in ACC reduced the performance accuracy (*y-axis*) when monkeys had lost more tokens in the near past (negative GTI) (random permutation, p<0.05). Data indicate the mean and the standard error of the mean. Accuracy on the trial-level is normalized by the mean and standard deviation of the accuracy in the baseline (the first six blocks prior to the TUS) in the same TUS sessions.

The slower learning in the gain-loss context after ACC-TUS was accompanied by prolonged explorations of objects prior to making a choice compared to pre-TUS baseline (Wilcoxon test, p=0.016), and compared to aSTR-TUS (Kruskal-Wallis test, p=0.036; pairwise Wilcoxon test, FDR multiple comparison corrected for dependent samples, p=0.03) (**Fig. 3A, Fig. S6A**,**C**). Explorative information sampling in the gain-only context did not vary between TUS conditions (Kruskal-Wallis test, p=0.55) (**Fig. 3B, Fig. S6B**,**D**). Once subjects reached the learning criterion in a learning context, they could exploit the learned feature rule until the block changed to a new feature rule after ∼30-55 trials. During this period, they showed overall high plateau performance which was significantly lower in the gain-loss (91%±0.07) than the gain-only contexts (92%±0.05; LME’s, t-stat= -3.95, p=0.001) (**Fig. S2C**). ACC-TUS exacerbated this performance drop in the gain-loss block, leading to significantly lower plateau accuracy compared to the pre-TUS baseline condition and compared to other TUS conditions in the gain-loss learning context (LME’s, t-stat= -2.05, p=0.04; **Fig. 3C**; **Fig. S6E**), but not in the gain-only learning context (Kruskal-Wallis test, p=0.8) (**Fig. 3D**; **Fig. S6F**).

We validated that the behavioral impairments emerged shortly after the sonication and lasted until the end of the ≤120 min. long session (**Fig. S7**). We also confirmed that the observed behavioral effects were not only evident when considering individual blocks (the block level analysis) (**Fig**.**s 1****-2**), but also when averaging the learning performance across blocks per session and applying session-level statistics (**Fig. S8, Supplementary Table 1**).

So far, we found that ACC-TUS slowed learning speed, increased exploration durations, and reduced plateau accuracy in the gain-loss contexts. These results could be due to motivational difficulties when adjusting to the experience of losing an already attained token. To analyze this adjustment to losses we looked at the performance in trials following choices that led to losses. We found that experiencing a loss leads to overall poorer performance in subsequent trials when TUS was directed to ACC but not after sham or aSTR-TUS (random permutation test, p<0.05, **Fig. 3E, Fig. S9A-D**). Importantly, this overall performance decrement was dependent on the recent history of losses. ACC-TUS reduced performance specifically when subjects had lost 2 or 3 tokens in the preceding four trials, but not when their net token gain in the past four trials was ≥0 tokens (random permutation test, p<0.05, **Fig. 3F**). This dependence of the ACC-TUS effect on the recent gross token income was evident in both monkeys (**Fig. S9E)** and could be a main source that led to slower learning.

## DISCUSSION

We found that sonicating the ACC, but not the anterior striatum, slowed down learning, prolonged visual information sampling of objects prior to choosing an object, and reduced overall performance when learning took place in a context with gains and losses and with high attentional load. These behavioral impairments were specific to the ACC-TUS condition when comparing behavior to baseline performance prior to TUS, and to other sham-controlled TUS and aSTR-TUS sessions. Moreover, the changes in behavioral adjustment were found when comparing average performance between individual sessions (**Fig. S8**), for performance variations across blocks of different sessions (**Fig. 2**), and on a trial-by-trial level in impaired behavior when the loss of tokens accumulated (**Fig. 3**). Taken together, these findings provide evidence that primate ACC is causally supporting the guidance of attention and information sampling in contexts that are motivationally challenging and cognitively demanding.

### Extending existing functional accounts of ACC functions

Our main findings suggest important extensions to two major theoretical accounts of the overarching function of the ACC for adaptive behavior. First, the finding of prolonged information sampling after ACC-TUS supports the view that the ACC is guiding information sampling and attention to maximize reward outcomes (*13, 22*). However, this functional deficit was only apparent in the gain-loss learning context in which subjects expected lower payoff, suggesting that the ACC guidance of information sampling particularly plays a role in motivationally challenging conditions. Secondly, our core finding of compromised learning in motivationally challenging conditions supports the view that ACC is essential to control effort (*14*). This functional impairment was evident when subjects faced both, high cognitive load and low payoffs for incorrect performance. High cognitive load alone was not sufficient to trigger an impairment. This result pattern is consistent with an effort control model that assumes the subjective costs of performance control is stronger influenced by the threat of loss than by experiencing high cognitive load. This finding is consistent with a wealth of data from affective processing, mostly from human imaging studies, that show systematic ACC activation in the face of aversive or threatening experiences (*23, 24*).

Taken together, the observed result pattern causally supports theories suggesting the ACC is essential for guiding information sampling as well as for controlling motivational effort during adaptive behavior. But our results suggest important constraints when the ACC is recruited for these functions. In particular, our results highlight that an intact ACC is essential to overcome motivational challenges and that this function is exacerbated when learning in a cognitively demanding situation. We discuss the specific rationale for this conclusion in the following.

### ACC modulates the efficiency of information sampling

We found that ACC-TUS increased the duration of fixating objects prior to choosing an object in the loss contexts. Longer durations of fixations are typically considered to reflect longer sampling of information from the foveated object. This has been inferred from longer fixational sampling of objects that have higher outcome uncertainty (*25*), or that explicitly carry information for reducing outcome uncertainty (*26, 27*). Consistent with this view, we found longer fixational sampling when objects had more feature information at high attentional load (**Fig. 2D**,**E**), and in the gain-loss context when subjects were uncertain not only about the gain outcome but additionally about the possible loss outcome (**Fig. 3D**,**E)**. ACC-TUS increased the duration of fixational sampling further in these conditions, particularly when subjects anticipated loss outcomes. This finding could be related to neurons in the intact ACC that fire stronger during the fixation of cues that reduces uncertainty about anticipated losses (*28*). In particular, trial-by-trial variations of ACC activity during the fixation of these cues has been found to correlate with trial-by-trial variations of seeking information about how aversive a pending outcome will be (*28*). Disrupting this type of activity with transcranial ultrasound in our study may thus have disrupted the efficiency of acquiring information while fixating objects. According to this interpretation, the TUS-induced prolongation of fixation durations indicates that an intact ACC causally enhances the efficiency of visual information sampling to reduce uncertainty about aversive outcomes.

### ACC mediates feature-specific credit assignment for aversive outcomes

The altered information sampling after ACC-TUS was only evident in the gain-loss context and depended on experiencing losing tokens. We found that the loss of one, two, or three tokens significantly impaired performance in ACC-TUS compared to sham sonication, or after striatal sonication (**Fig. 3F**). This valence-specific finding might be linked to the relevance of the ACC for processing negative events and using negative experiences to adjust behavior. Imaging studies have consistently shown ACC activation in response to negatively valenced stimuli or events (*23, 24, 29*). Behaviorally, the experience of loss outcomes is known to trigger autonomous vigilance responses that reorient attention and increases explorative behaviors (*16, 30, 31*). Such loss-induced exploration can help avoid threatening stimuli, but it comes at the cost of reducing the processing depth for the loss-inducing stimulus <u>(Banaie Boroujeni et al., 2020)</u>. Studies in humans have shown that stimuli associated with loss of monetary rewards or aversive outcomes (aversive images, odors, or electrical shocks) are less precisely memorized than stimulus features associated with positive outcomes (*32*–*34*). This effect is sometimes considered to reflect the over-generalization of aversive outcomes to stimuli that only share some resemblance with the precise loss-inducing stimuli. Such over-generalization is evolutionary meaningful because it allows fast recognition of a stimulus as potentially threat-related even if it is not a precise instance of a previously encountered, loss-inducing stimulus <u>(Banaie Boroujeni et al., 2020; Moscarello and Hartley, 2017)</u>. For our task, such a precise recognition of features of a chosen object was pivotal to learn from feature-specific outcomes. Our finding of reduced learning from loss outcomes therefore indicates that ACC-TUS exacerbated the difficulty to assign negative outcomes to features of the chosen object.

Neurophysiological support for this interpretation of the ACC to mediate negative credit assignment comes from studies that report activity changes of the ACC neurons to reward prediction error was selectively encoding the specific object feature that gave rise to the unexpectedly negative (loss) outcome (*36*). This study also showed that feature-specific prediction error signals in the ACC predict when neurons update value expectations about specific features when encountering those features in the future <u>(Oemisch et al., 2019; aee also: Quilodran et al., 2008)</u>. Thus, neurons in ACC signal feature-specific negative prediction errors and the updating of feature-specific value predictions. It seems likely that disrupting these signals with TUS might have led to impaired feature-specific credit assignment in our task. This scenario is supported by the finding that ACC-TUS impaired flexible learning most prominently at high load, i.e. when there was a high degree of uncertainty about which stimulus features were distracting features associated with loss and which was the target feature associated with gain (**Fig. 2C**). This finding suggests that outcome processes such as credit assignment in an intact ACC causally contribute to the learning about aversive distractors.

### A role of the ACC to determine learning rates for aversive outcomes

So far, we have discussed that ACC-TUS has likely reduced the efficiency of information sampling and this reduction could originate from the disruption of feature-specific credit assignment processes. These phenomena were exclusively observed in the loss context, and ACC-TUS did not change learning or information sampling in the gain-only context. Based on this finding, we propose that the ACC plays a causal role in determining the learning rate for negative or aversive outcomes and thereby controls how fast subjects learn which object features in their environments have aversive consequences. Such learning from negative outcomes was particularly important for our task at higher load (*38*), because at high load, a single loss outcome provided unequivocal information that all features of the chosen object were non-rewarded distracting features. This was different for positive outcomes which were ambiguous about which of the two or three features of the chosen object was causing the positive outcome. The higher informativeness of negative than positive outcomes can explain why ACC-TUS caused a selective learning impairment in the loss context. This interpretation is consistent with the established role of the ACC to encode different types of errors (*39*–*41*) and with computational evidence that learning from errors and other negative is dependent on a dedicated learning mechanism separately of learning from positive outcomes (*38, 42*–*44*).

The suggested role of the ACC to determine the rate of learning from negative outcomes describes a mechanism for establishing which visual objects are distractors and should be actively avoided or actively suppressed in a given learning context (*45*). When subjects experience aversive outcomes, the ACC may use these experiences to bias attention and information sampling away from the negatively associated stimuli. This suggestion is consistent with rapid-onset activity changes of the ACC neurons to the onset of covert attention cues that trigger the inhibition of processing distracting and enhancement of processing target information (*46, 47*). In these studies, the rapid activity modulations reflect a push-pull attention effect that occurred during covert attentional orienting and was independent of actual motor actions. This observation supports the general notion that ACC circuits are critical for guiding covert and overt information sampling during adaptive behaviors (*13*).

### Versatility of the ultrasound protocol for transcranial neuromodulation

Our conclusions were made possible by using a transcranial ultrasound stimulation protocol that was developed to interfere with local neural activity in deep neural structures and impose a temporary functional disconnection of the sonicated area from larger networks (*18, 18, 20*). We implemented an enhanced protocol that quantified the high anatomical targeting precision (**Fig. S3A**) and confirmed that sonication power reached the targets (**Fig. S3B-E**). We also documented that the main ACC-TUS behavioral effect (impaired learning) was evident relative to a within-task pre-sonication baseline and throughout the experimental behavioral session (**Fig. S7**). Importantly, the main behavioral effects of ACC-TUS in our task are consistent with the effects from more widespread, invasive lesions of the ACC in non-human primates. Widespread ACC lesions have resulted in gradual reductions of affective response to harmful stimuli (*48*), response perseverations (*49*), failures to use reward history to guide choices (*7, 8*) and reduced control to inhibit prevalent motor programs (*50, 51*). Our study therefore illustrates the versatility of the transcranial ultrasound stimulation approach to disrupt deeper structures such as the ACC that so far have been out of reach for noninvasive neuromodulation techniques such as transcranial magnetic stimulation (TMS) or transcranial direct current stimulation (tDCS) (*52*).

In summary, the pattern of our results illustrates that the ACC multiplexes motivational effort control and attentional control functions by tracking the costs of incorrect performance, by optimizing feature-specific credit assignment for aversive outcomes, and by actively guiding information sampling to visual objects during adaptive behaviors.

## Acknowledgments

This work was supported by the National Institute of Mental Health of the National Institutes of Health under Award Number R01MH123687 and 1UF1NS107666. The authors thank Dr. Marcus Watson for help with the experimental software, Huiwen Luo for assistance with calibrating optically tracked devices using MRI and Adrienne Hawkes for technical assistance with voltage monitoring. The content is solely the responsibility of the authors and does not necessarily represent the official views of the National Institutes of Health.

## Author Contributions

K.B.B., C.F.C. and T.W. conceived the experiments. K.B.B. and R.L.T. performed the experiment. M.K.S. and T.M. contributed the image-guided procedures and computer simulations. K.B.B. reconstructed the brain images. K.B.B. and M.K.S. wrote the code to run the experiments. K.B.B. analyzed and visualized the data. All authors contributed writing the paper.

## Data and code availability

All data supporting this study and its findings, as well as custom MATLAB code generated for analyses, are available from the corresponding author upon reasonable request.

## Fig.s and Fig. Legends

**Fig. S1.**
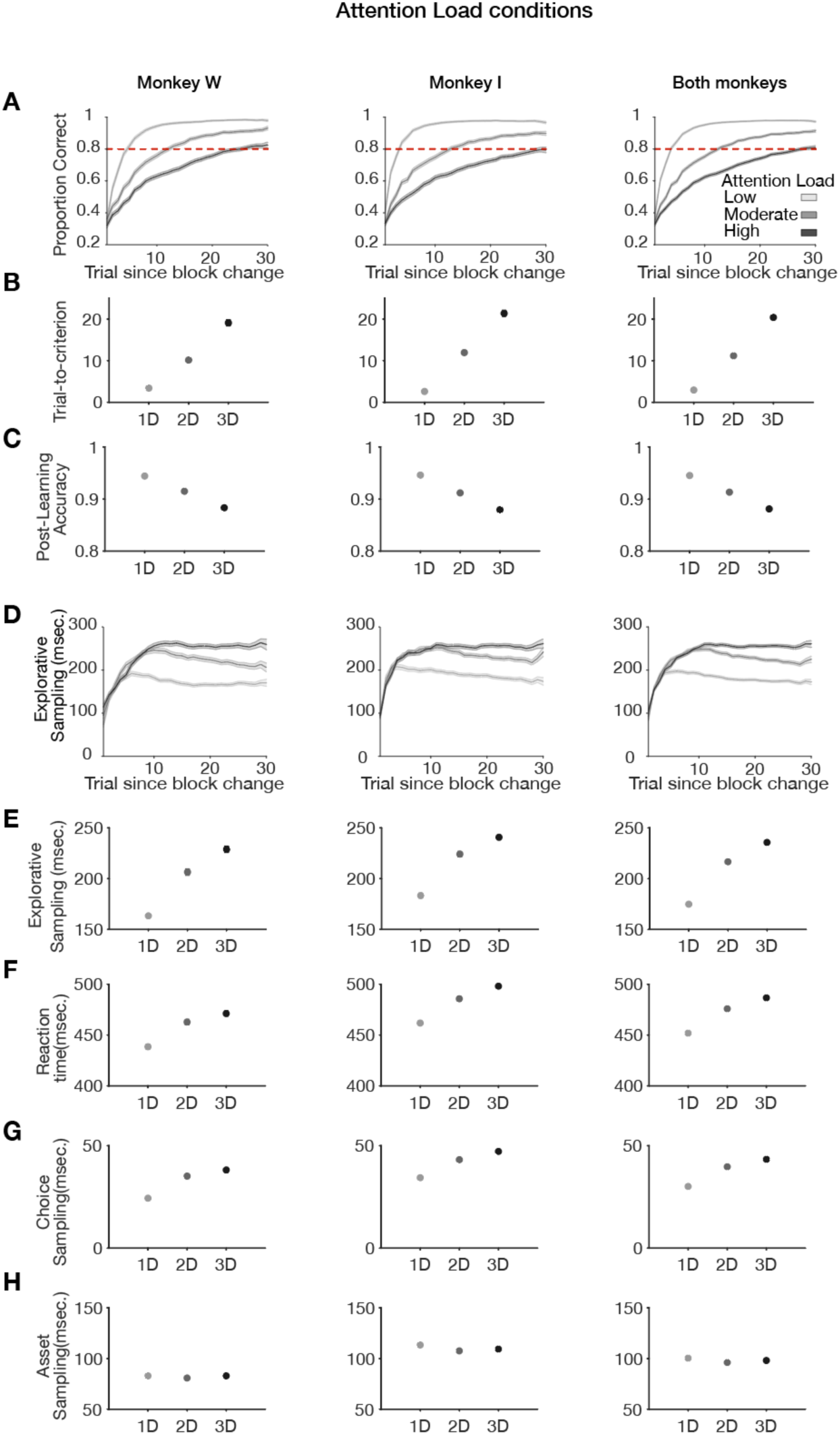
Attentional load effect on learning and fixational information sampling. **A**,**B**. Both monkeys reached the learning criterion of 80% or more correct trials (based on a 12 trials forward-looking window). Learning is fastest at low attentional load (*light grey*), and slowest at high attentional load (*dark grey*). In all panels, the left column shows the results for monkey W, the middle for monkey I, and the right for both monkeys combined. **C**. Post-learning accuracy is significantly reduced in higher attentional load (LME’s, P<0.001). **D**,**E**. Explorative sampling is the duration in msec. of fixational sampling of objects prior to making a choice. Explorative sampling increased at the beginning of a block, reached a maximum during learning, and but remained elevated only at the highest attentional load (LMEs, p<0.001). **F**. Choice reaction time is the time from stimulus onset to the onset of the final fixation (that chooses the object). It increased with attentional load (Kruskal-Wallis test, p<0.001). **G**. Choice sampling is measured as the duration of sampling the chosen object prior to the final choice fixation. Choice sampling is increased in blocks with higher attentional load (Kruskal-Wallis test, p<0.001). **H**. Asset sampling duration quantifies how long subjects fixate the token bar prior to choosing an object. Asset sampling is independent of attentional load, and more extensive in monkey I. Data show means and standard error of the mean.

**Fig. S2.**
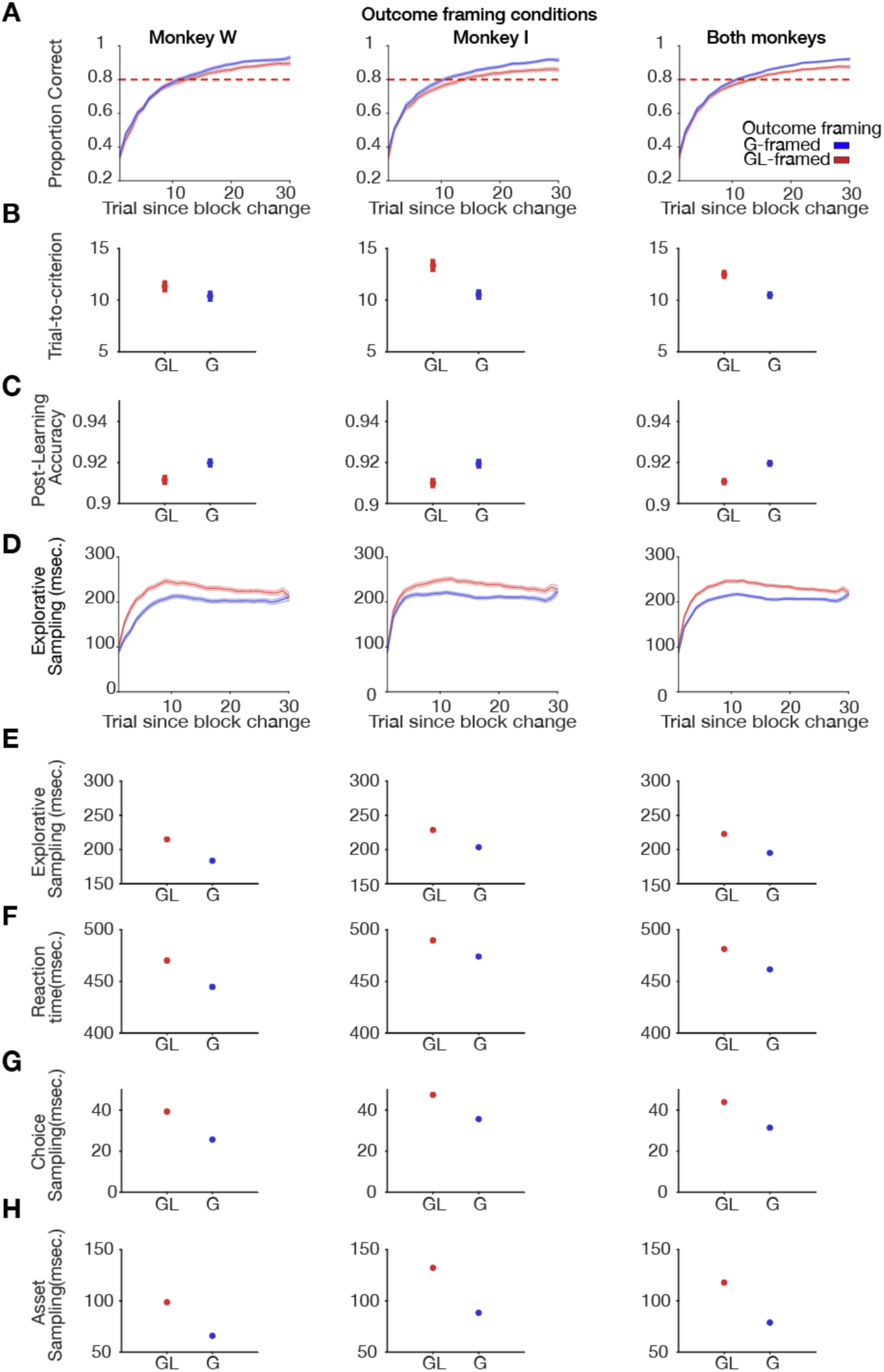
Effects of Gain-Loss and Gain-Only learning contexts on learning and gaze sampling behaviors. Same format as **Fig. S1. A**,**B**. Learning is faster in the gain-only context (blue), than in the gain-loss context. In all panels, the left column shows the results for monkey W, the middle for the monkey I, and the right for both monkeys. **C**. Post-learning accuracy is significantly reduced in the gain-loss context (Wilcoxon test, P<0.001). **D**,**E**. Explorative sampling reaches a higher maximum during learning in the gain-loss context (red) compared to the gain-only context (blue) (Wilcoxon test, P<0.001). **F**. Choice reaction time is slower in the gain-loss context (Wilcoxon test, p<0.001). **G**. Sampling of the object that is subsequently chosen is longer in the gain-loss context (Wilcoxon test, p<0.001). **H**. Asset sampling duration is longer in the gain-loss context (Wilcoxon test, p<0.001). Data show mean and standard error of the mean.

**Fig. S3.**
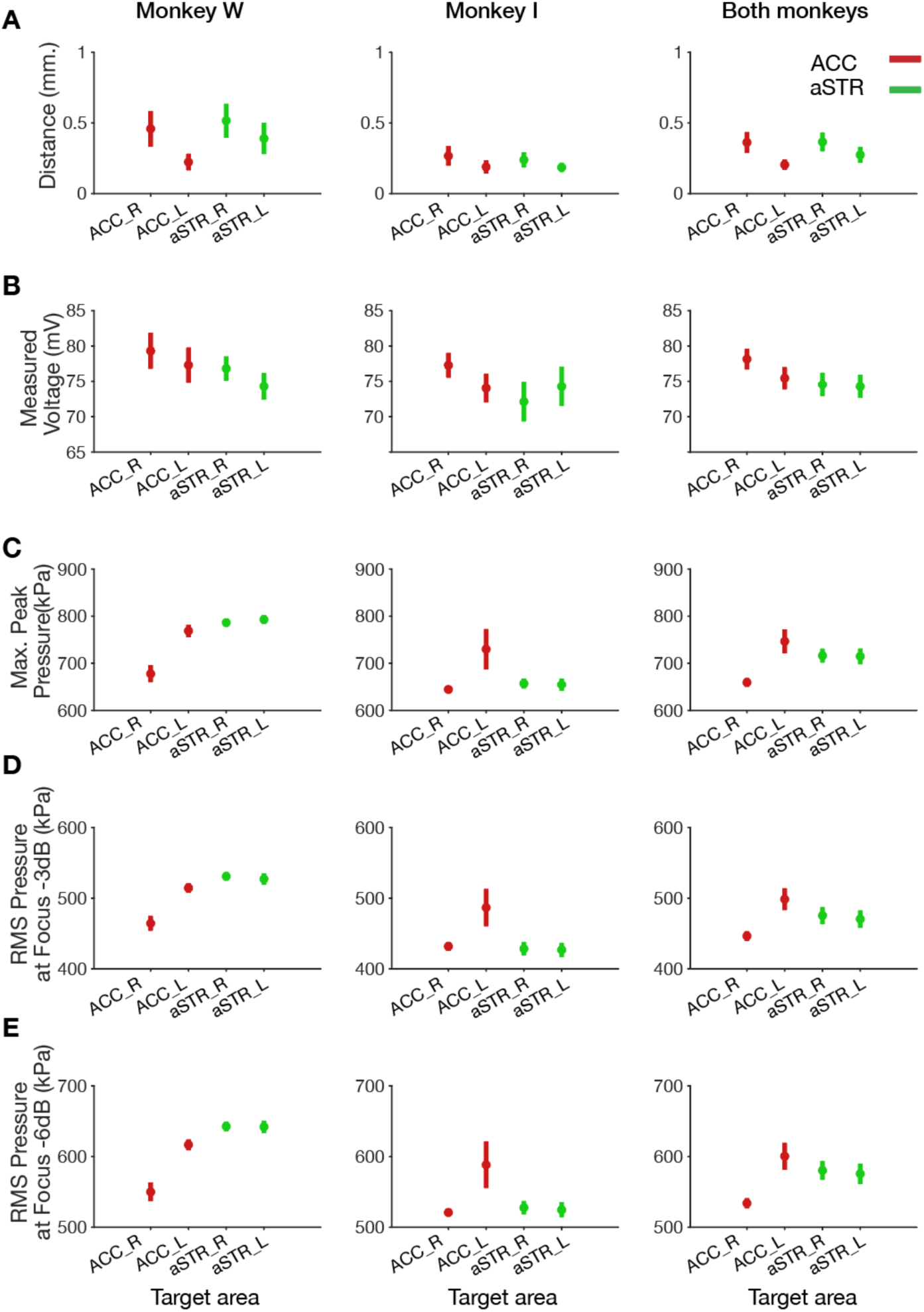
Transcranial ultrasound stimulation localization, energy, and sonication focus specifications. In each experimental session we positioned the transducer sonication beam to focus on left and right hemisphere ACC and striatum and sonicated the area for 40 seconds in each hemisphere. **A**. Both hemispheres (ACC in red, and anterior striatum in green) in both monkeys were targeted precisely within an averaged sub-millimeter distance of the center of focal beam of the transducer to the anatomical target region. **B**. Real-time monitoring of output voltage to the transducer confirmed reliable feedforward voltage range for both monkeys (STAR Methods). **C**. Computer simulations show a reliable range of maximum peak pressure at the focal center of the transducer. **D**,**E**. Root mean squared (RMS) deviation values of negative peak pressure at the focus of the sonication attenuated at -3dB (**D**), and -6dB (**E**).

**Fig. S4.**
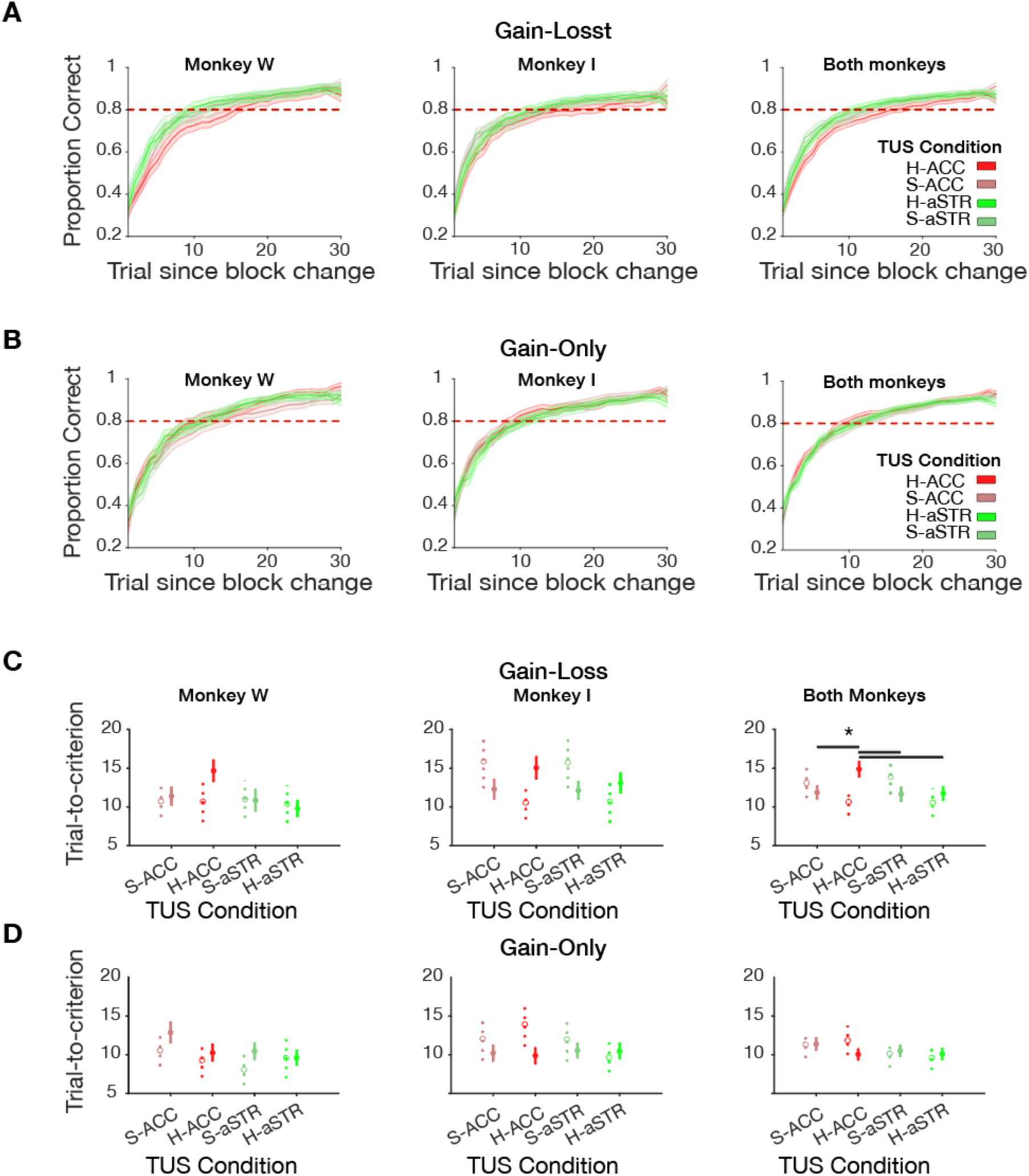
Transcranial ultrasound stimulation effect on learning for individual monkeys (left and middle column) and their average (right column). **A**,**B**. Learning curves for four different experimental conditions: high energy TUS in ACC (H-ACC; red), or anterior striatum (H-aSTR; green), or sham ACC (S-ACC; darkened red), or sham anterior striatum (S-aSTR; darkened green). Learning curves are shallower after ACC-TUS in the gain-loss context (**A**) but not in the gain-only context (**B**). **C**,**D**. The reduced learning speed (increased trials-to-criterion) with TUS-ACC in the gain-loss learning context (**C**) but not in the gain-only learning context (**D**). Detailed statistics are provided in **Supplementary Table 1**. Data show mean and standard error of the mean.

**Fig. S5.**
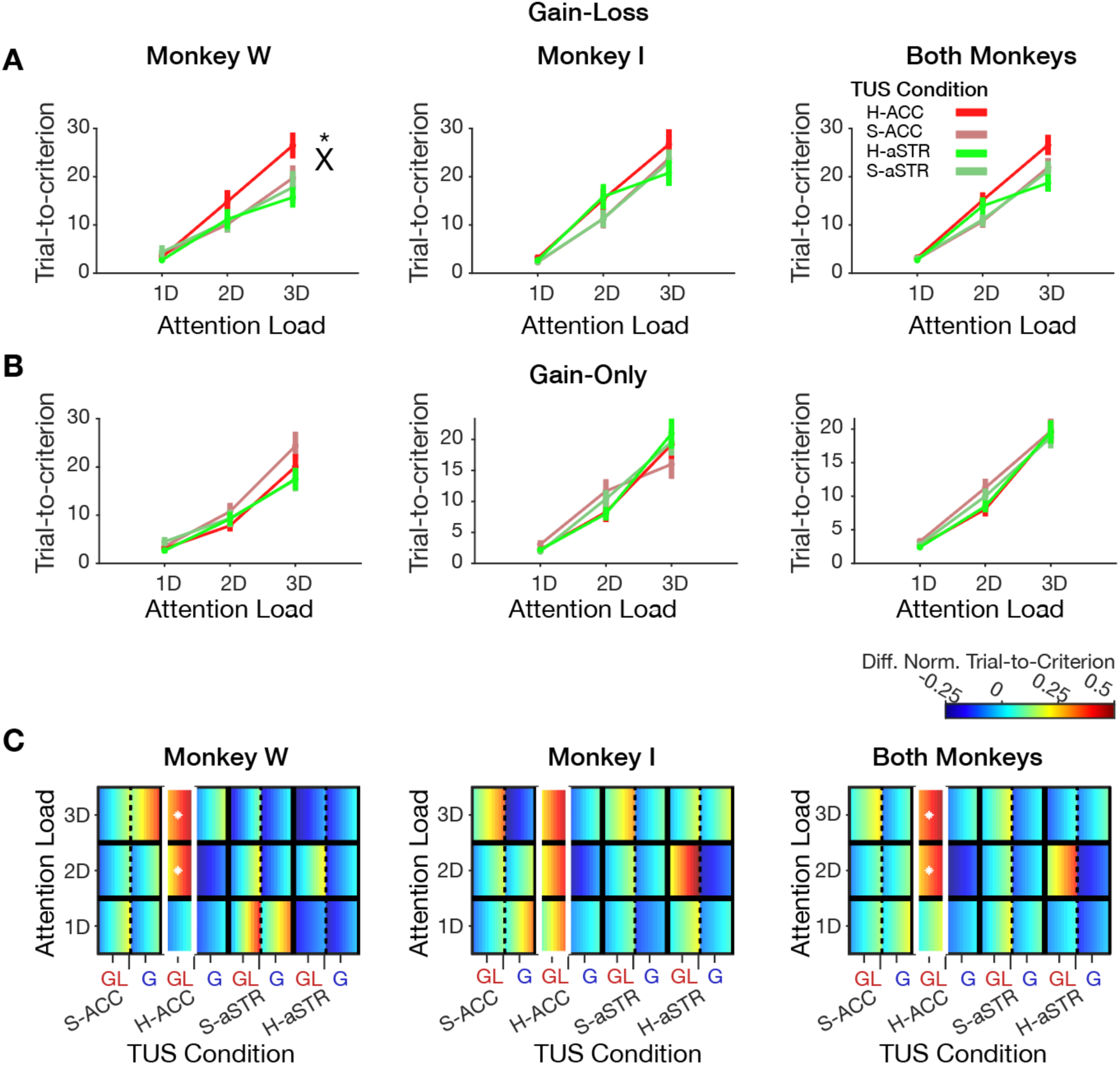
TUS interaction of attentional load and motivational context. **A**,**B**. TUS in ACC reduces learning speed at higher attentional load in the gain-loss learning context (**A**), but not the gain-only learning context (**B**) (for the full multiple comparison corrected statistical results see **Supplementary Table 2**) **C**. Marginally normalized trials-to-criterion are significantly higher with ACC-TUS in the gain-loss learning context, and the effect in ACC-TUS was only significant at higher attentional load conditions 2D and 3D (random permutation p<0.05). Error bars are standard error of the mean. The white rectangle in (**C**) shows learning in a TUS condition is different from other TUS conditions. Each cell is color coded with the mean value ± standard error of the mean with a low to high value gradient from left to right. The white asterisk shows attentional loads in a learning context in a TUS condition is significantly different from other TUS conditions. In all panels, the left column shows the results for monkey W, the middle for the monkey I, and the right for both monkeys combined.

**Fig. S6.**
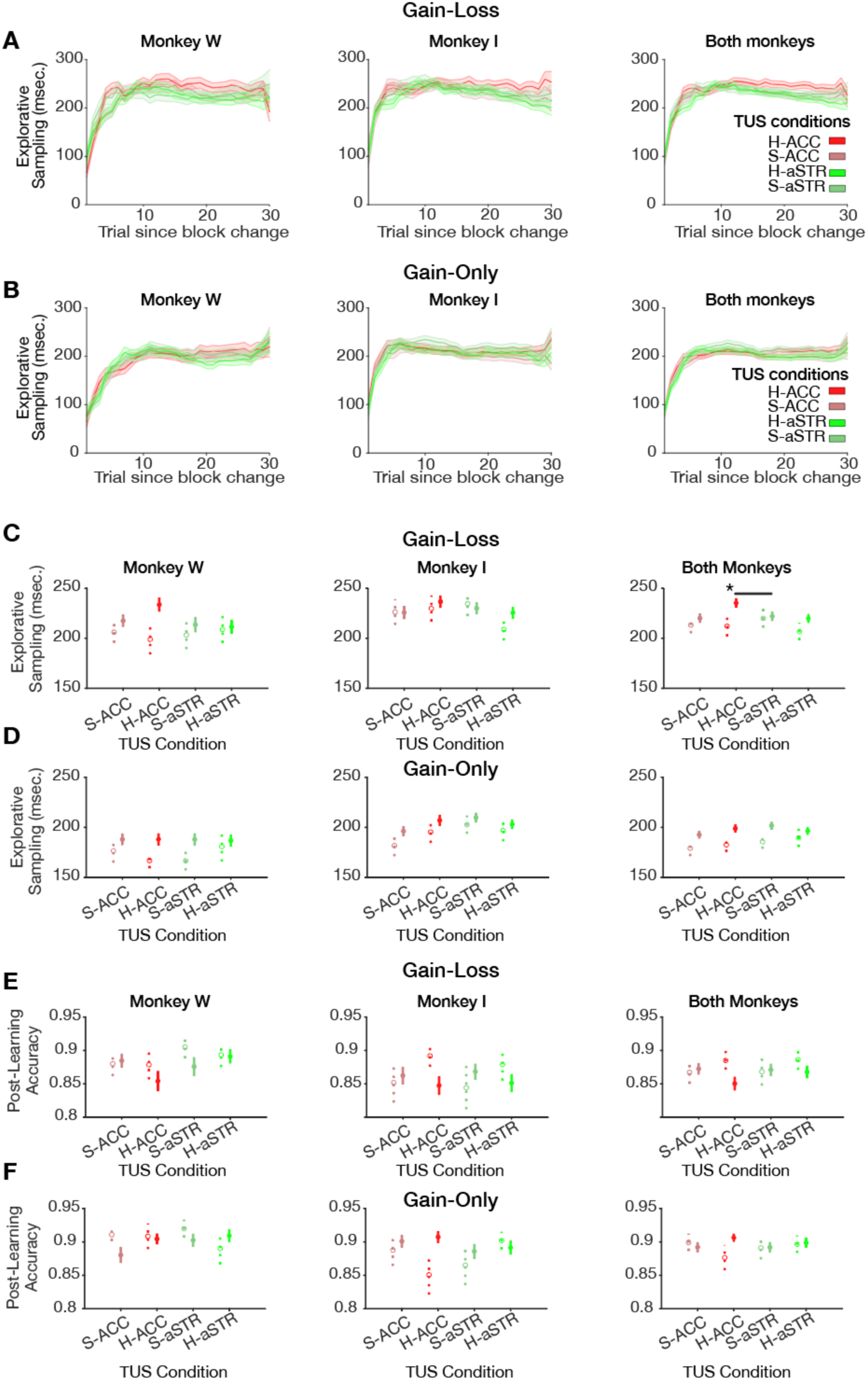
Transcranial ultrasound stimulation effects on explorative behavior. **A**,**B**. Explorative sampling curves showing the average fixation durations on objects before making a choice. With the beginning of a new learning block explorative sampling increases to a maximum during learning and then slowly reduces to a baseline level. TUS in ACC in the gain-loss learning context (**A**), but not in the gain-only context causes elevated exploratory sampling (**B**). **C**,**D**. Explorative sampling is increased with TUS in ACC relative to baseline and other TUS conditions in the gain-loss context (**C**) but not the gain-only context (**D**). **E**,**F**. Post-learning accuracy is not significantly different from the baseline and other TUS conditions in the block-level analysis, neither in gain-loss context (**E**), nor the gain-only context (**F**) (detailed statistics in **Supplementary Table 1)**.

**Fig. S7.**
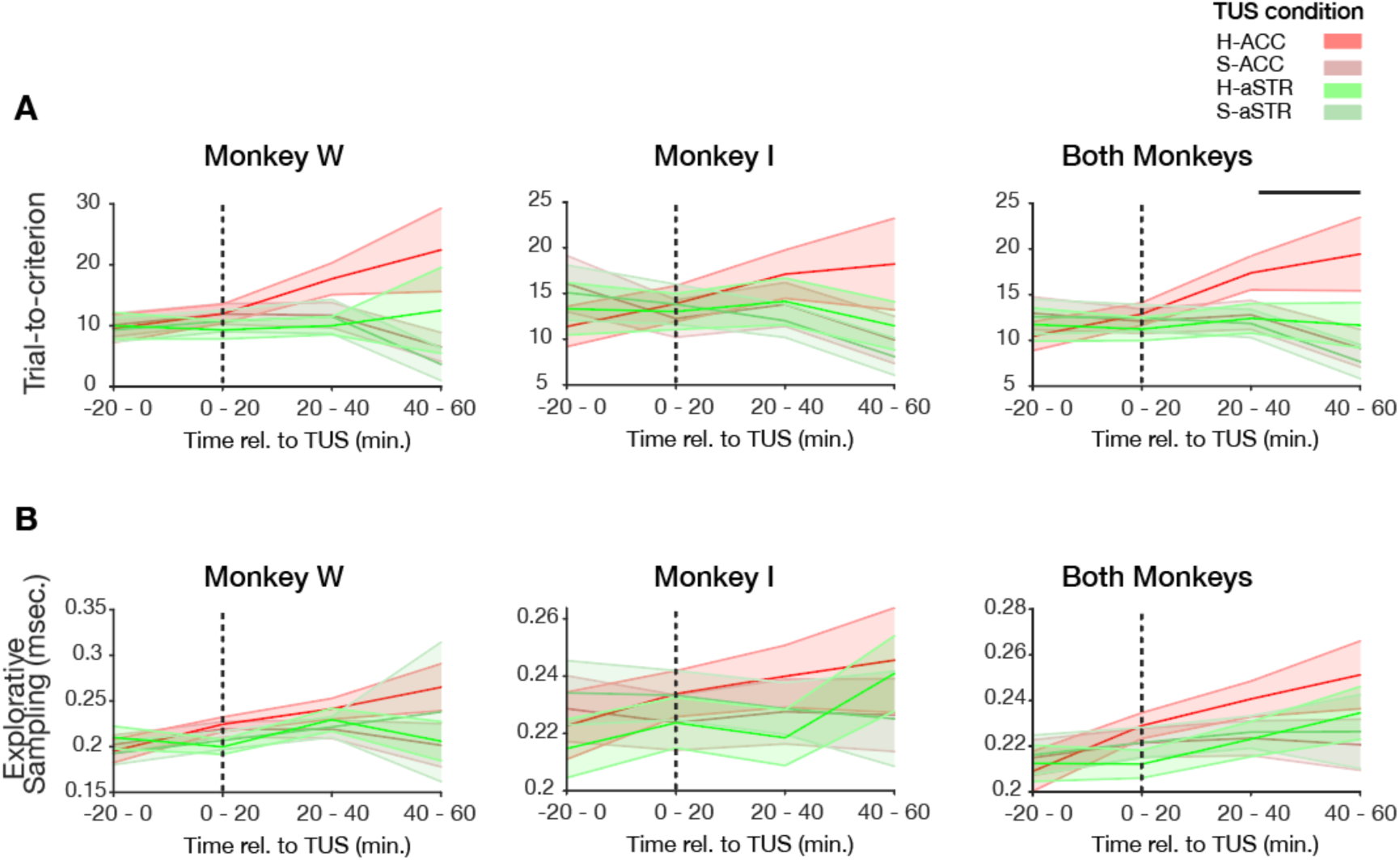
Time course of the effect of transcranial ultrasound stimulation on learning and explorative behavior. **A**. Monkeys show a gradual deterioration of learning with TUS in ACC over time in the gain-loss conditions. **B**. Explorative sampling increases over time relative to the time of TUS in ACC. The x-axis shows the time in minutes relative to TUS stimulation within a session. The time-step resolution is 20 minutes. Both monkeys show a similar time course with the slowing of learning from 20-40 minutes after the stimulation to the end of the session (randomized permutation test, p<0.05). Shaded error bars are standard errors of the mean.

**Fig. S8.**
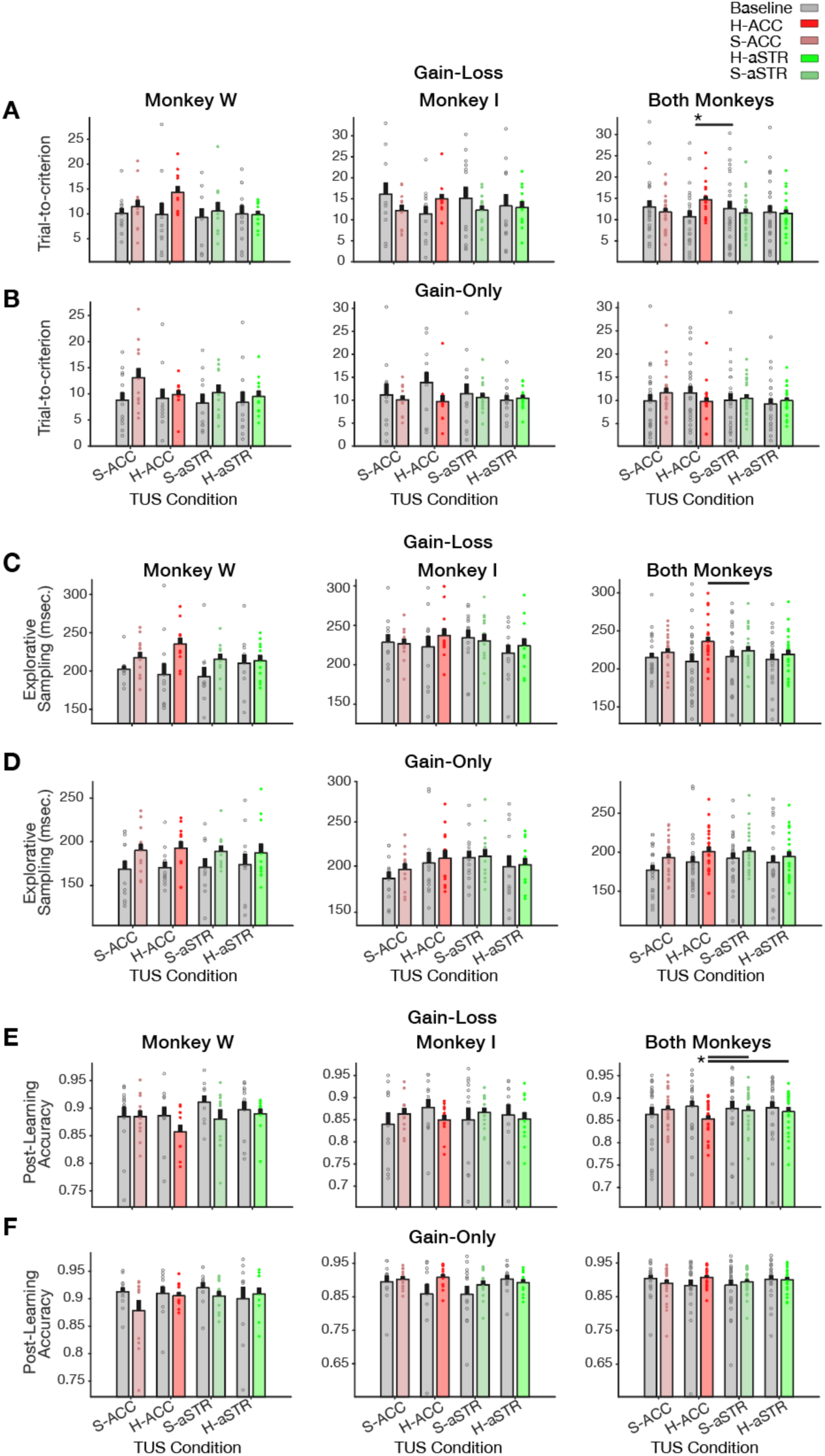
Session level effect of transcranial ultrasound stimulation on learning and explorative sampling. Average results for each session and across sessions for monkey W and I, and both combined (left, middle, and right column, respectively). **A**,**B**. TUS in ACC caused slower learning in both monkeys in the gain-loss context (**A**), but not in the gain-only context (**B**). **C**,**D**. TUS in ACC causes longer explorative sampling than Sham aSTR in the gain-loss context (**C**). There was, however, no difference to baseline or in the gain-only context. (**D**). **E**,**F**. TUS in ACC reduced post-learning accuracy relative to baseline and other TUS conditions in the gain-loss context (**E**), but not the gain-only context (**F**). For detailed statistics see **Supplementary Table 1**. Data show session means for the pre-stimulation baseline blocks (*grey*) and across blocks after TUS/Sham (*colored*). Significant pairwise comparisons are indicated with black horizontal lines between each pair of conditions and the black asterisk shows significant session-level difference (post TUS versus baseline) of each behavioral measure.

**Fig. S9.**
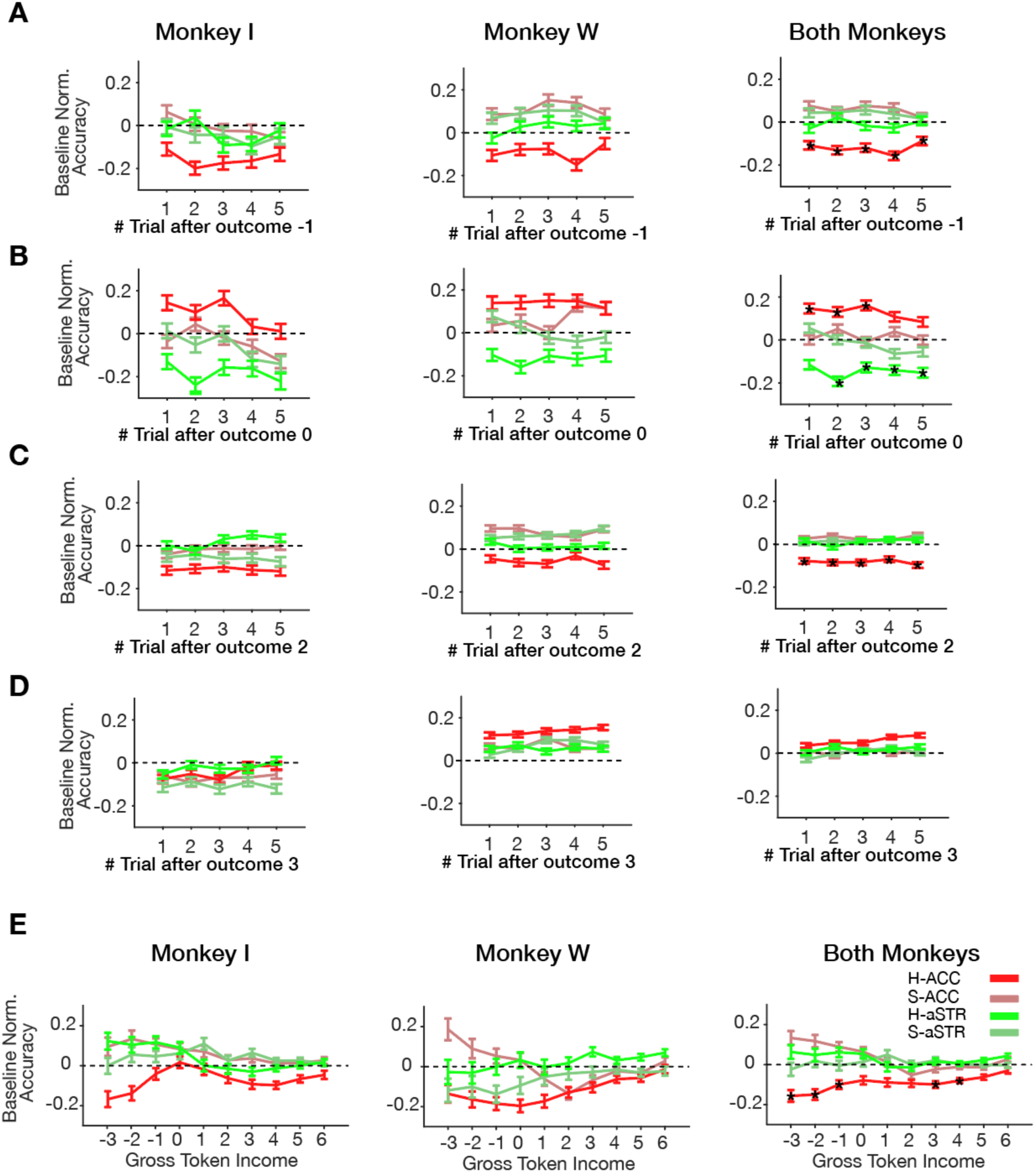
Trial-level effects of transcranial ultrasound stimulation on post-outcome performance adjustment. **A**-**D**. Normalized performance accuracy in the five trials after receiving either a loss of one token (**A**), no loss after an incorrect response (**b**), a gain of 2 tokens (**C**), and a gain of 3 tokens (**D**). TUS in ACC (**red**) caused overall reduced performance accuracy in each monkey after losing 1 token or gaining 2 tokens which shows the effect is specific to the gain-loss context. **e**. Both monkeys show reduced performance (y-axis) after TUS in ACC when the Gross token income (GTI) was negative, i.e. then they on average had lost 1-3 tokens in the preceding five trials. Accuracy on the trial-level analysis is normalized by the mean and standard deviation of the accuracy of similar events in the baseline during the same TUS session. All statistics in these panels used randomization (permutation) tests with FDR correction of p-values for dependent samples with an alpha-level of 0.05. The black asterisk shows points significantly different from pre-TUS baseline and relative to other TUS conditions.

**Supplementary Table 1.**
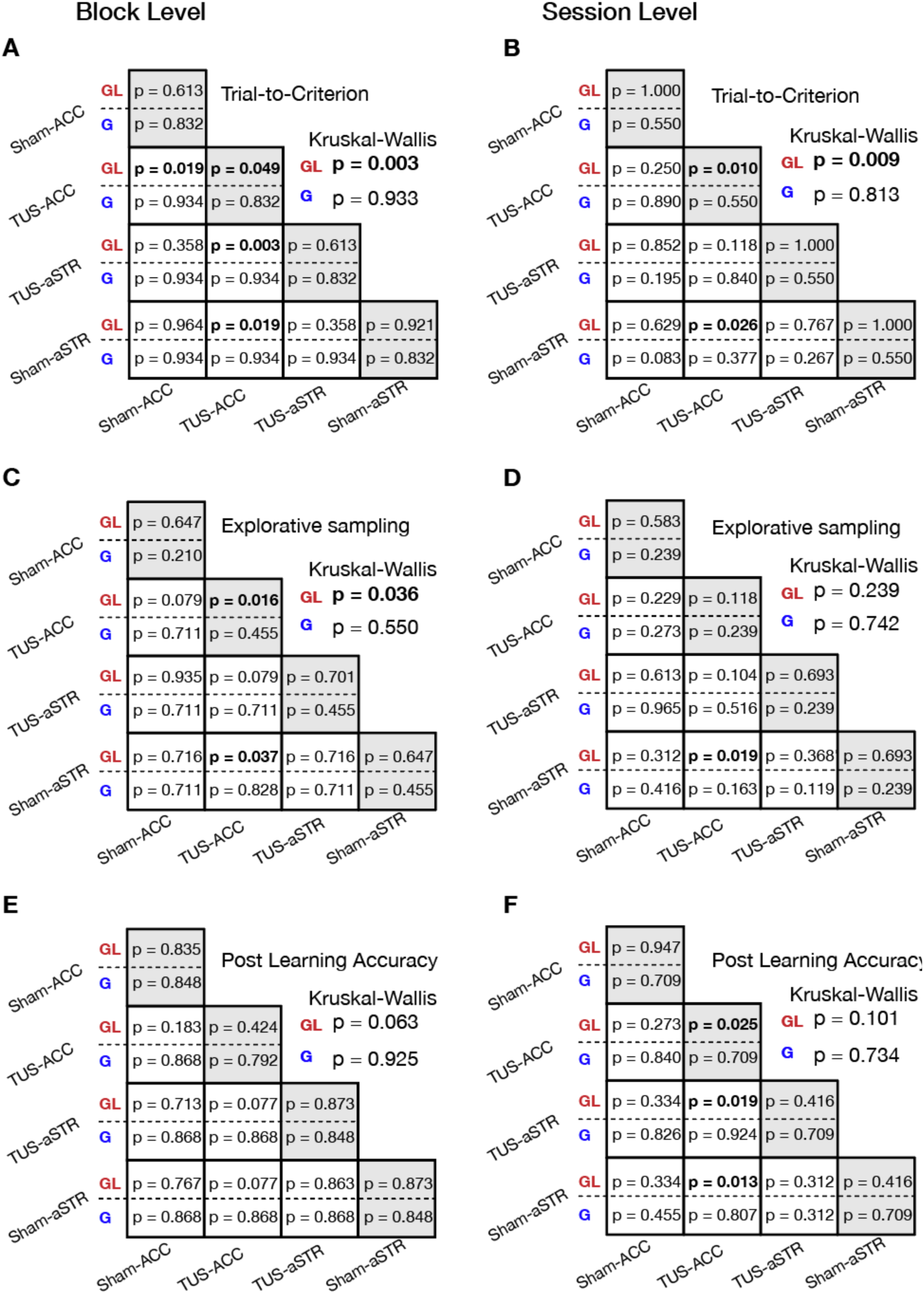
Statistical results considering data from individual blocks (block level, left) and average data from individual sessions (session level: right) for trials-to-criterion (**A**,**B**), explorative sampling durations (**C**,**D**), and the post-learning plateau accuracy (**E**,**F**). Each table shows results of 3 different tests for the 2 motivational learning contexts: gain-only (G, in blue font), and gain-loss (GL, in red font). The p-values on the diagonal show Wilcoxon tests for each TUS condition compared to its baseline (before the stimulation). The non-diagonal cells show p-values for pairwise comparisons of each pair of the TUS conditions. The overall Kruskal-Wallis test results is shown on the right outside of each table. All p-values are FDR-corrected with an alpha level of 0.05, as explained in the methods.

**Supplementary Table 2.**
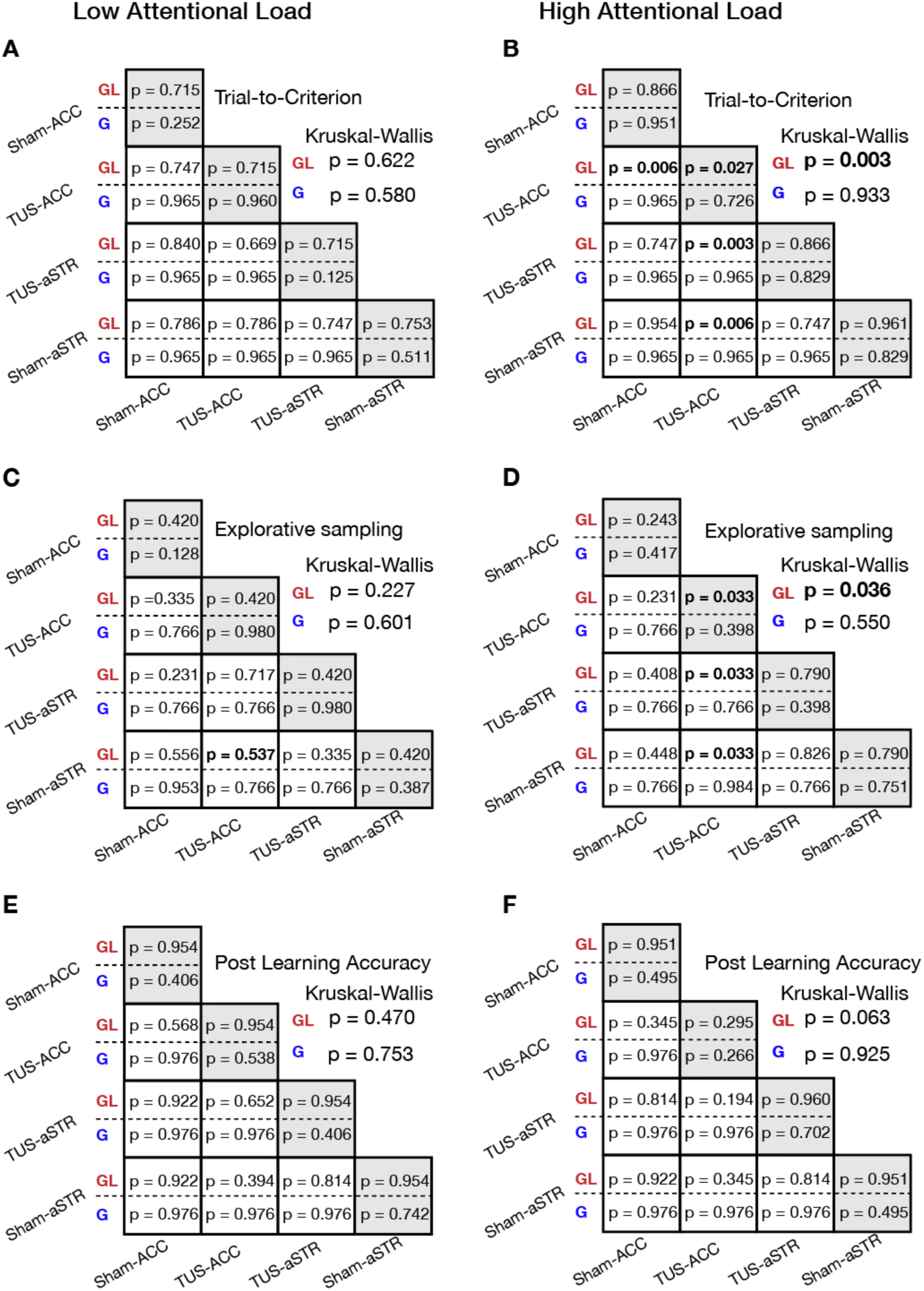
Statistical results split for low (left) and high (right) attentional load conditions for trials-to-criterion (**A**,**B**), explorative sampling durations (**C**,**D**), and the post-learning plateau accuracy (**E**,**F**). Each table shows results of 3 different tests for the 2 motivational learning contexts: gain-only (G, in blue font), and gain-loss (GL, in red font). The p-values on the diagonal show Wilcoxon tests for each TUS condition compared to its baseline (before the stimulation). The non-diagonal cells show p-values for pairwise comparisons of each pair of the TUS conditions. The overall Kruskal-Wallis test results is shown on the right outside of each table. All p-values are FDR-corrected with an alpha level of 0.05, as explained in the methods.

## MATERIALS AND METHODS

### Experimental procedures

All procedures were in accordance with the National Institutes of Health Guide for the Care and Use of Laboratory Animals, the Society for Neuroscience Guidelines and Policies, and approved by the Vanderbilt University Institutional Animal Care and Use Committee. Two male macaque monkeys (monkey I 13.6 kg, and monkey W 14.6 kg, 8-9 years of age) contributed to the experiments. They sat in a sound-proof booth in primate chairs with their head position fixed, facing a 21’’ LCD screen at a distance of 63 cm from their eyes to the screen center. Behavior, visual display, stimulus timing, and reward delivery were controlled by the *Unified Suite for Experiments* (USE), which integrates an IO-controller board with a unity3D video-engine based control for displaying visual stimuli, controlling behavioral responses, and triggering reward delivery (*53*). Prior to the ultrasound experiment, the animals were trained on the feature learning task in a kiosk training station (*54*). Monkeys first learned to choose objects to receive an immediate fluid reward before a token system that provided animals with green circles per correct choice that symbolized tokens later cashed out for fluid reward. Tokens were presented above the chosen object and traveled to a token bar where they accumulated with successive correct performance. When five tokens were collected, the token bar blinked red/white, fluid was delivered through a sipper tube, and the token bar reset to five empty token placeholders (**Fig. 1A**). The animals effortlessly adopted the token reward system as documented in <u>(Banaie Boroujeni et al., 2020)</u>. Here, we used in separate blocks of 35-50 trials a condition with ‘gains-only’ (3 tokens for correct choices, no penalties) and with ‘gains-and-losses’ (2 tokens for correct choices and 1 token lost, i.e., removed from the token bar, for incorrect choices). The introduction of gains-and-losses effectively changed behavior. Animals learned slower, showed reduced plateau accuracy, enhanced exploratory sampling and more checking of the token bar (**Fig. S2B-H**).

### Task paradigm

The task required monkeys to learn feature-reward rules in blocks of 35-60 trials through trial-and-error by choosing one of three objects. Objects were composed of multiple features, but only one feature was associated with reward (**Fig. 1A-C**). A trial started with the appearance of a black circle with a diameter of 1 cm (0.5° radius wide) on a uniform grey background on the screen. Monkeys fixated the black circle for 150 ms to start a trial. Within 500 ms after the central gaze fixation registration, three objects appeared on the screen randomly at 3 out of 4 possible locations with an equal distance from the screen center (10.5 cm, 5° eccentricity). Each stimulus had a diameter of 3 cm (∼1.5° radius wide). To choose an object, monkeys had to maintain fixation onto the object for at least 700 ms. Monkeys then had 5 sec. to choose one of three objects or the trial was aborted. Choosing the correct object was followed by a yellow halo around the stimulus as visual feedback (500 ms), an auditory tone, and either 2 or 3 tokens (green circles) added to the token bar (**Fig. 1A**). Choosing an object without the rewarded target feature was followed by a blue halo around the selected objects, a low-pitched auditory feedback, and in the loss conditions, the presentation of a grey ‘loss’ token that traveled to the token bar where one already attained token was removed. The timing of the feedback was identical for all types of feedback. In each session, monkeys were presented with up to 36 separate learning blocks, each with a unique feature-reward rule. Across all 48 experimental sessions, monkeys completed on average 23.6 (±4 SE) learning blocks per session (monkey W: 20.5±4, monkey I: 26.5±4). For each experimental session, a unique set of objects was defined by randomly selecting three dimensions and three feature values per dimension (e.g., 3 body shapes: oblong, pyramidal and ellipsoid; 3 arm types: upward pointy, straight blunt, downward flared; 3 patterns: checkerboard, horizontal striped, vertical sinusoidal; <u>(Watson et al.,</u> have documented the complete library of features). Of this feature set three different task conditions were defined: One task condition contained objects that varied in only one feature while all other features were identical, i.e., the object body shapes were oblong, pyramidal, and ellipsoid, but all objects had blunt straight arms and uniform grey color. A second task condition defined objects that varied in two feature dimensions (‘2D’ condition), and a third task condition defined objects that varied in three feature dimensions (‘3D’ condition). Learning is systematically more demanding with increasing number of feature dimensions that could contain the rewarded feature (for computational analysis of the task, see <u>Womelsdorf et al., 2021)</u>. We refer to the variations of the object feature dimensionality as attentional load because it corresponds to the size of the feature space subjects have to search to find the rewarded feature (**Fig. 1E, Fig. S1**). 1D, 2D, and 3D conditions varied randomly across blocks.

### Experimental design

Each session randomly varied across blocks two motivational conditions (Gain/Loss and Gains-only) and three attentional load conditions (1D, 2D, 3D). Randomization ensured that all (six) combinations of conditions were present in the first six blocks prior to sonication and that all combinations of conditions were equally often shown in the 24 blocks after the first six blocks. After monkeys completed the first 6 learning blocks (on average 12 minutes after starting the experiment), the task was paused to apply transcranial ultrasound stimulation. After bi-lateral placement of the transducer and sonication (or sham sonication) of the brain areas, the task resumed for 18-24 more blocks (**Fig. 1D**). Experimental sessions lasted 90-120 minutes.

### Transcranial Ultrasound Stimulation (TUS)

For transcranial stimulation, we used a single element transducer with a curvature of 63.2 mm and an active diameter of 64 mm (H115MR, Sonic Concepts, Bothell, WA). The transducer was attached to a cone with a custom-made trackable arm. Before each session, we filled the transducer cone with warm water and sealed the cone with a latex membrane. A conductive ultrasound gel was used for coupling between the transducer cone and the shaved monkey’s head. A digital function generator (Keysight 33500B series, Santa Rosa, CA) was used to generate a periodic burst of 30 ms stimulation with a resonate frequency of 250 kHz and an interval of 100-ms for a total duration of 40-second and 1.2 MPa pressure (similar to (*19, 56*)) (**Fig. 1D**). A digital function generator was connected to a 150-Watt amplifier with a gain of 55 dB in the continuous range to deliver the input voltage to the transducer (E&I Ltd., Rochester, NY). We measured the transducer output in water using a calibrated ceramic needle hydrophone (HNC 0400, Onda Corp., Sunnyvale CA) and created a linear relationship between the input voltage and peak pressure. To avoid hydrophone damage, only pressures below a mechanical index (MI) of 1.0 were measured, and amplitudes above this were extrapolated. We have previously measured transducer output at MI>1.0 with a calibrated optical hydrophone (Precision Acoustics, Dorchester, UK) to validate the linearity of this relationship at higher MI, but this calibrated device was not available during these studies. During stimulation, the bi-directionally coupled (ZABDC50-150HP+, Mini Circuits Brooklyn, NY) feedforward and feedback voltage were monitored and logged using a Picoscope 5000 series (A-API; Pico Technology, Tyler, TX) and a custom written python script.

Four different sonication conditions were pseudo-randomly assigned to the experimental days per week for a 12-week experimental protocol per monkey. These four conditions consisted of high energy TUS in anterior striatum (H-aSTR), high energy TUS in anterior cingulate cortex (H-ACC), sham anterior striatum (S-aSTR), and sham anterior cingulate cortex (S-ACC) (**Fig. 1D**). We sequentially targeted an area in both hemispheres (each hemisphere for a 40 sec duration) with real-time monitoring of the distance of the transducer to the targeted area (**Fig. S3A**) and monitoring of the feedforward power (**Fig. S3B**). Sham conditions were identical to TUS conditions, only no power was transmitted to the transducer.

### Neuro-navigation setup

An MR image was acquired before implantation of the head-post and a CT image was obtained containing the headpost with a fiducial array arm attached to it during the scan. The fiducial array in the CT scan was identical to its position during the neuro-navigation procedure in an experimental session. We co-registered MR and CT images into the same space (CT images were the fixed volume to prevent distortion of fiducial array space) using FSL (*57*). 3D-Slicer software (http://www.slicer.org), with Image-guided therapy (IGT) (*58*) extension modules were used for tracking and positioning the transducer. First, the co-registered brain images were imported to 3D-Slicer. Fiducial array points were marked on the images ordered from right to left. These points allowed reconstructing the relative distances of the sonication target locations in the brain from the fiducial array. At the beginning of each session, the fiducial array was mounted to monkey’s headpost and fixed to the exact location as in the CT images. A stylus tool was then used to collect the physical fiducial array points in the same order as it was marked in the image space. After collecting the physical points, we transformed the physical space into image space. The focus to tracker transformation (which was used for tracking the focus of the transducer relative to the tracker attached to its enclosing cone) was also transformed with the same transformation to the image space (*59, 60*). We then navigated the focal point of the transducer (a circle with 3 mm radius) centered at the maximum focus estimated as sphere model (1-mm radius) placed at the focal maximum that was used for the location of targets in each area and hemisphere (these sphere models were held the same across all experiments for each monkey). Under the IGT module, we tracked the real-time distance of the center of the transducer focus from the target sphere model.

### Acoustic modeling of sonication beam

To validate sonication of the ACC and aSTR, we performed 3D numerical simulations of the acoustic propagation through the monkey skull using the open-source MATLAB toolbox k-Wave (Http://www.k-wave.org). A single-element transducer with a radius of curvature of 63.2 mm and an active diameter of 64 mm (H115MR, Sonic Concepts, Bothell, WA) and resonant frequency of 250 kHz was modeled as a spherical section. We used 3D-Slicer combined with optical tracking to position the transducer for simulations to recreate the optical tracking geometry. To localize the transducer relative to the transducer tracker, we manually aligned a transducer mask’s marked geometric focus created in k-Wave with the center of the focal location from reconstructed temperature images collected using a 2D gradient echo thermometry pulse. MR thermometry was performed on a 3.0T Philips Achieva 2D gradient echo thermometry pulse sequence with a multi-shot EPI readout factor of 3. Imaging parameters were 150 × 150 mm^2^ field of view (FOV), 112 × 112 matrix, 1.3 × 1.3 mm^2^ voxel size, 5 slices, 4 mm slice thickness, TE/TR 12/500 msec. A continuous wave sonication for 30 sec. at input voltage of approximately 1 MPa was used to generate several degrees of heating. This calibration was performed once to generate a transform with an accurate offset and rotation relative to the optical tracker. A CT image of the respective monkey was resampled using the Resample Image (BRAINS) module in Slicer so that the resolution and spacing matched the calibrated transducer mask or integration with simulations. The resampled data was exported to MATLAB to run acoustic simulations, using the CT to estimate the speed of the sound and density of the skull. The final spatial maps were imported back into 3D Slicer and overlaid for comparison with the estimated focus from the optical tracking and MR thermometry results. To account for heterogeneities of the monkey skull, a linear relationship between Hounsfield units (HU) from CT scans was used to calculate the speed of sound and density (*61*). Brain tissue was assumed to have the same speed of sound and density as water. Values used included a padded grid size [Nx,Ny,Nz] of [384,288,280] with isometric voxels of 0.25 mm, attenuation = 8 dB/cm, power law absorption exponent of 1.1, *c*_*min*_ = 1480 *m/s, c*_*max*_ = 3100 *m/s, ρ*_*water*_ = 1000 *kg/m*^3^, and *ρ*_*bone*_ = 2100 *kg/m*^3^. Maximum pressure was recorded for every voxel in the simulation grid. The highest pressure was found at the skull, as it absorbs more sound than brain tissue. For each dataset, the spatial map cropped the maximum pressure of the skull so that an accurate root-mean-square (RMS) of the sound pressure could be calculated at the focus. The focal spot size of the sonicated area was calculated using *P* > *P*_*max*_/2 for half-maximum pressure (−3 dB; **Fig. S3D**) and 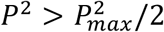 for half maximum intensity (−6 dB; **Fig. S3E**, error bars showing standard error of the mean across sessions). The half-maximum pressure and intensity equations were used to create a mask of the focal spot volume and calculate the RMS pressure for each (**Fig. 2A**).

### Analysis of behavioral learning and fixational sampling

The improvement of accuracy over successive trials relative to the beginning of a learning block reflects the learning curve, which we computed with a forward-looking 12-trial averaging window in each block (**Fig. 1E**). We defined learning speed in a block as the number of trials needed to reach criterion accuracy of ≥80% correct trials over the subsequent 12 trials (**Fig. 2B**,**C**; **Fig. S1C, Fig. S2C**). When monkeys did not reach the learning criterion at the end of a block, we estimated the trial number the monkeys would have learned the block with a linear regression fit to the performance accuracy of monkeys in the last 12 trials in the block. We computed the plateau performance accuracy, or *post-learning accuracy*, as the proportion of correct choices across all trials after the learning criterion was reached in a block (**Fig. 3C**,**D**). In blocks in which the learning criterion was not reached, we computed the accuracy over the last 12 trials in the block. Analysis of exploratory eye movements used an adaptive velocity-based thresholding algorithm (*62, 63*) to detect saccadic eye movements and fixations (*i*) onto objects during explorative sampling prior to choosing an object (*explorative sampling*), (*ii*) onto the chosen object prior to choosing it (*choice fixations*), and (*iii*) onto the token bar (*asset sampling*) (see e.g., **Fig. S1E**,**G**,**H** and **Fig. S2E**,**G**,**H**).

### Trial-level statistical analysis

We tested TUS effects on behavior at the trial level using linear mixed effect models (LME) (*64*) with 4 main factors: *attentional load (Att*_*Load*_) with three levels (1D, 2D, and 3D distractor feature dimensions, ratio scale with values 1,2 and 3), *trial in block (TIB), previous trial outcome* (*Prev*_*Outc)*_) which is the number of tokens gained or lost in the previous trial, motivational token condition, which we call the *motivational Gain/Loss context (MCtx*_*Gain/Loss*_) with two levels (1, for the loss condition, and 2 for the gain condition, nominal variable), TUS condition (*TUS*_*Cnd*_) with four levels (Sham-ACC, High-ACC, Sham-STR, High-STR), and *time relative to stim (T2Stim)* with 2 levels (before versus after stimulation). We used 3 other factors as random effects, a factor *target features* (*Feat*) with 4 levels (color, pattern, arm, and shape), weekday of the experiment (*Day*) with 4 levels (Tuesday, Wednesday, Thursday, and Friday), and the factor *monkeys* with 2 levels (W and I). We used these factors to predict 3 metrics (*Metric*): accuracy (*Accuracy*), reaction time (*RT*), explorative sampling (*Sample*_*Explr*_). The LME is formalized as in eq. 1.

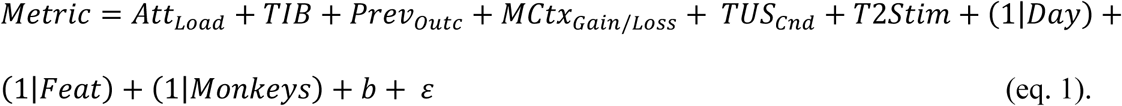

### Analysis of behavioral adjustment to the recent token history (Gross Token Income)

To quantify how TUS affected choice accuracy depending on the motivational status of the subjects, we calculated the gross sum of the earned token over four trials which we call *gross token income* (GTI; **Fig. 3F, Fig. S9E**). For each value of the GTI (spanning from – 3 to +6 tokens), we calculated the accuracy on the subsequent trial and normalized it by subtracting the mean and dividing by the standard deviation of the same GTI values during the baseline. We then asked whether the accuracy of choice for a given GTI was changed relative to baseline or other TUS conditions in any of the TUS conditions. We first used Wilcoxon signed-rank test to test for each TUS condition and each GTI, whether accuracy changes relative to baseline were different from zero. We then used FDR control of multiple comparison for dependent samples (*65*), with an alpha level of 0.05 to adjust p-values across different GTIs and the TUS conditions. In a second analysis, we applied randomization statistics to test whether, for a given GTI, the accuracy change was different compared to TUS conditions. We randomly permuted trial labels of the TUS conditions and selected 1000 subsamples with a sample size equal to the true TUS and GTI conditions. We then formed a randomized sampling distribution of the means and extracted the probability of randomly finding values more extreme than the true baseline-corrected accuracy value for each TUS and GTI condition, under the null hypothesis that for a given GTI none of the TUS conditions show accuracy changes relative to the baseline different than other TUS conditions. After finding the p-values for all GTI conditions, we corrected them for multiple comparisons using FDR control at an alpha level of 0.05. GTI values for a TUS condition were considered significant if it passed both tests (**Fig. 3F, Fig. S9E**).

### Analysis of previous-trial-outcome effects on accuracy adjustment

We measured whether TUS modulated the accuracy on the following 1-5 trials after experiencing gain or loss outcomes (**Fig. 3E, Fig. S9A-D**). Accuracy was baseline normalized for each TUS condition. Wilcoxon rank tests were applied to test the effects of the previous trial outcome after versus before TUS with FDR correction for dependent samples (*65*) with an alpha level of 0.05. In a next step, we tested post-outcome accuracy changes in one TUS condition compared to other TUS conditions. Thus, for each outcome condition, we randomly permuted the TUS condition label of the trials, and randomly selected 1000 subsamples with a sample size equal to the size of the true outcome and the TUS condition (both in here and in the GTI analysis, only the TUS labels were shuffled and the rest of the labels for trials e.g., the order relative to TUS was still the same). We then used the 1000 subsamples and formed a randomized sampling distribution of the means and calculated the probability of finding a value more extreme than the true mean of changed trials accuracy following an outcome relative to the baseline, under the null hypothesis that the accuracy changes relative to the baseline after a given trial outcome is not different across different TUS conditions. After finding the p-values for each TUS condition and n^th^ (n=1-5) following trials, we used FDR correction for dependent samples with an alpha level of 0.05 to adjust p-values across different post outcome trials and the TUS conditions.

### Block-level analysis of behavioral metrics

We used linear mixed effect models to analyze across blocks how learning speed (indexed as the ‘learning trial’ (*LT*) at which criterion performance was reached), post-learning accuracy (*Accuracy*), and metrics for explorative and choice sampling were affected by TUS. In addition to the fixed and random effects factors used for the LME in eq. 1 we also included the factor block *switching condition* (*Switch*_*Cnd*_) with 2 levels (intra- and extra-dimensional switch). The LME had the form of eq. 2:

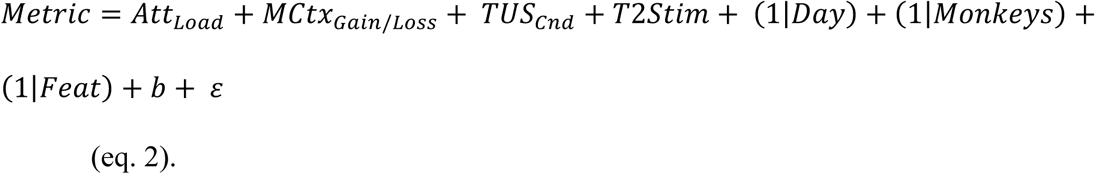

We extended the model to test for interactions of *TUS*_*Cnd*_, *MCtx*_*Gain/Loss*_, and *T*2*Stim*:

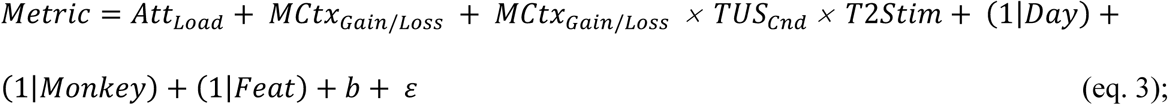

for interactions of *T*2*Stim, Att*_*Load*_ and *TUS*_*Cnd*_.:

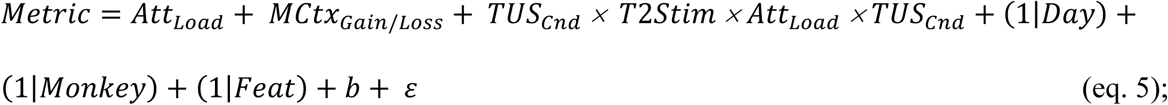

and for interactions of *T*2*Stim, Att*_*Load*_, *MCtx*_*Gain/Loss*_ and *TUS*_*Cnd*_.:

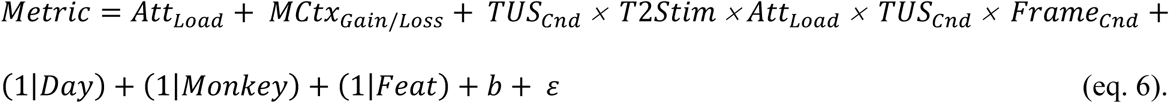

Where appropriate we used a non-parametric approach by fitting generalized linear mixed effects models (GLME’s) as in equations 2-6. While the formulas were the same as in LME’s, we used a link *Identity* function, and a *Poisson* distribution for the response variable.

### Pairwise comparisons of TUS effects

As a validation of the overall mixed effect modeling results, we report a Kruskal Wallis test for a main effect of TUS conditions on behavioral metrics and follow up with Wilcoxon tests for all pairwise comparisons between individual TUS conditions at each motivational context (gains-only and gains-loss). P-values were FDR corrected to control for multiple comparisons of dependent samples (*65*). Complete statistical results are provided in **Supplementary Table 1**. We repeated the same procedure for pairwise comparisons of TUS effects at low attentional load (1D condition) and high attentional load (combining 2D and 3D conditions) (**Supplementary Table 2**).

### Randomization statistics for normalized behavioral metrics

We calculated the normalized value of a behavioral metric across blocks in each TUS condition, motivational contexts, and attentional loads by subtracting their marginal means and dividing by the standard deviation. As shown in **Fig. 2D** and **Fig. S5C** this procedure provides a table where each cell shows the mean of the marginally normalized metric value ± the standard error of normalized metric value (for each cell, the color shows from left to right the mean minus the standard error to the mean plus the standard error of that cell). We then performed randomization tests at two levels. First, at the level of the motivational contexts, we asked whether any TUS condition significantly differed from others by permuting the TUS condition labels and randomly selecting 1000 subsamples with a size equal to the size of the true TUS condition. This tested the null hypothesis that the marginally normalized metric values in a given TUS condition in a given motivational context do not differ from other TUS conditions in the same motivational context. We calculated the p-values and adjusted them post-hoc using FDR correction for dependent samples with an alpha level of 0.05. The white rectangles in **Fig. 2D** and **Fig. S5C** show the FDR corrected, significant TUS. In a second analysis, we applied the same rationale and procedure but this time the randomization test was done on each motivational context and attentional load condition separately to test the null hypothesis that TUS conditions had no effect on the behavioral metrics. Those TUS conditions that showed a significant difference from others in a given attentional load and motivational context are marked with a white asterisk in **Fig. 2D** and **Fig. S5C**.

### Session-Level analysis of TUS effects on behavioral metrics

In addition to analyzing behavior in individual blocks and performing block-level statistics (see above), we also analyzed the data across sessions and provide these session-level results in **Supplementary Table 1** and **Fig. S8**. For each session, we calculated the mean of each behavioral metric for gain-only and gain-loss learning contexts separately before and after the TUS. We then used Wilcoxon tests for each TUS condition under the null hypothesis that the behavioral metric is not different before versus after TUS. To compare behavioral metrics across TUS conditions we first applied a Kruskal-Wallis test under the null hypotheses that the behavioral metrics did not differ between TUS conditions. Then we used pairwise Wilcoxon comparisons to compare each pair of TUS conditions. All p-values were corrected post-hoc using FDR for dependent samples with an alpha level of 0.05

